# Distinct groups of RNA viruses associated with thermoacidophilic bacteria

**DOI:** 10.1101/2023.07.02.547447

**Authors:** Syun-ichi Urayama, Akihito Fukudome, Miho Hirai, Tomoyo Okumura, Yosuke Nishimura, Yoshihiro Takaki, Norio Kurosawa, Eugene V. Koonin, Mart Krupovic, Takuro Nunoura

**Affiliations:** Department of Life and Environmental Sciences, Laboratory of Fungal Interaction and Molecular Biology, University of Tsukuba, 1-1-1 Tennodai, Tsukuba, Ibaraki 305-8577, Japan; Microbiology Research Center for Sustainability (MiCS), University of Tsukuba, 1-1-1 Tennodai, Tsukuba, Ibaraki 305-8577, Japan; Howard Hughes Medical Institute, Department of Biology and Department of Molecular and Cellular Biochemistry, Indiana Univeristy, Bloomington, IN, USA; Super-cutting-edge Grand and Advanced Research (SUGAR) Program, Japan Agency for Marine Science and Technology (JAMSTEC), 2–15 Natsushima-cho, Yokosuka, Kanagawa 237– 0061, Japan; Marine Core Research Institute, Kochi University, 200 Otsu, Monobe, Nankoku City, Kochi, 783-8502, Japan; Research Center for Bioscience and Nanoscience (CeBN), JAMSTEC, 2–15 Natsushima-cho, Yokosuka, Kanagawa 237–0061, Japan; Department of Science and Engineering for Sustainable Innovation, Faculty of Science and Engineering, Soka University, Hachioji 192-8577, Japan; National Center for Biotechnology Information, National Library of Medicine, Bethesda, MD, USA; Institut Pasteur, Université Paris Cité, CNRS UMR6047, Archaeal Virology Unit, Paris, France

**Author notes:** Corresponding author: Name: Syun-ichi Urayama.

## Abstract

Recent massive metatranscriptome mining substantially expanded the diversity of the bacterial RNA virome, suggesting that additional groups of riboviruses infecting bacterial hosts remain to be discovered. We employed full length double-stranded (ds) RNA sequencing for identification of riboviruses associated with microbial consortia dominated by bacteria and archaea in acidic hot springs in Japan. Whole sequences of two groups of multisegmented riboviruses genomes were obtained. One group, which we denoted hot spring riboviruses (HsRV), consists of unusual viruses with distinct RNA-dependent RNA polymerases (RdRPs) that seem to be intermediates between typical ribovirus RdRPs and viral reverse transcriptases. We also identified viruses encoding HsRV-like RdRPs in moderate aquatic environments, including marine water, river sediments and salt marsh, indicating that this previously overlooked ribovirus group is not restricted to the extreme ecosystem. The HsRV-like viruses are candidates for a distinct phylum or even kingdom within the viral realm *Riboviria*. The second group, denoted hot spring partiti-like viruses (HsPV), is a distinct branch within the family *Partitiviridae*. All genome segments in both these groups of viruses display the organization typical of bacterial riboviruses, where multiple open reading frames encoding individual proteins are preceded by ribosome-binding sites. Together with the identification in bacteria-dominated habitats, this genome architecture indicates that riboviruses of these distinct groups infect thermoacidophilic bacterial hosts.

## Introduction

During the last decade, metagenomics and metatranscriptomics analyses have transformed the study of viromes. These approaches that do not require laborious virus cultivation have become the principal source of virus discovery as reflected in the formal recognition of the sufficiency of virus genome sequences assembled from metagenomes and metatranscriptomes as prototypes of virus species or higher taxa ^1^. Indeed, numerous virus groups across all taxonomic levels have been discovered. In particular, the diversity of RNA viruses that, in the current comprehensive virus taxonomy, comprise the kingdom *Orthornavirae* within the realm *Riboviria* expanded dramatically through global metatranscriptome analyses ^2-6^. Three recent extensive metatranscriptome surveys led to the increase of number of identified ribovirus species by more than an order of magnitude ^7-9^.

Only one hallmark gene encoding the RNA-dependent RNA polymerase (RdRP) is conserved across the entire kingdom *Orthornavirae*. Therefore, detection of the RdRP, typically, using sensitive search methods based on sequence profiles, is the principal approach employed in metatranscriptome mining for riboviruses, and phylogenetic analysis of the RdRP is the basis of ribovirus taxonomy. Prior to the advent of massive metatranscriptome analysis, the viruses in this kingdom have been classified into 5 large phyla corresponding to major clades in the RdRP phylogeny ^10^. Metatranscriptome studies largely validated the robustness of these phyla although some groups of viruses have been reallocated between the phyla. In addition, several candidate smaller phyla were identified whereas the diversity of riboviruses across the lower taxonomy ranks, from class down to species demonstrated a nearly uniform increase, for example, roughly, five-fold, in one study that provided quantitative estimates ^8^. Apart from the major increase in the knowledge on RNA virus diversity, metatranscriptome mining for riboviruses yielded qualitative insights into the global view of the RNA virome. Traditionally, riboviruses have been recognized as the major component of the eukaryote virome whereas the viromes of bacteria and archaea were dominated by DNA viruses ^11,12^. For many years, only two small families of RNA viruses each infecting a narrow range of bacteria have been known, *Leviviridae* (single-stranded RNA bacteriophages) and *Cystoviridae* (double-stranded RNA bacteriophages). Metatranscriptome analyses revealed a much greater diversity of leviviruses than previously suspected, elevating this family to rank of the class *Leviviricetes* that includes multiple family and order level groups, in particular, the most abundant family of riboviruses, *Steitzviridae* ^8,13-15^. The family *Cystoviridae* was substantially expanded as well ^8^. When it comes to uncharacterized groups of viruses without a close relationship to any known ones, host assignment becomes a challenge. Nevertheless, several converging lines of evidence including (nearly) exclusive co-occurrence with bacteria, prediction of multiple virus genes preceded by prokaryote-type (Shine-Dalgarno) ribosome-binding sequences (RBS), identification of virus-encoded cell wall lysis enzymes, and perhaps most notably, targeting by reverse transcriptase (RT) containing type III CRISPR systems strongly suggest that several previously uncharacterized groups of riboviruses infect prokaryotes ^8^. At least two of these groups showed no close relationships with any of the 5 previously identified phyla and are candidates for new phyla. Others comprise the order *Durnavirales* within the phylum *Pisuviricota* that unites dsRNA viruses of families *Cystoviridae* and *Picobirnaviridae*, several uncharacterized families, and a subset of the expansive family *Partitiviridae*. Both *Picobirnaviridae* and *Partitiviridae* have been previously associated with eukaryotic hosts, but the results of metatranscriptome analysis call for this assignment to be reassessed. More generally, these findings indicate that the diversity of riboviruses infecting bacteria has been substantially underestimated and that additional groups of such viruses remain to be discovered.

Long dsRNA is a molecular marker of RNA virus infection ^16^. The recently developed method of Fragmented and primer-Ligated DsRNA Sequencing (FLDS) made it possible to capitalize on the presence of (nearly) identical terminal sequences in genome segments of the same virus. This information enables us to identify multi-segmented RNA virus-like genomes even if they did not show sequence similarity to known viruses ^17-19^. Here, we used FLDS for identification of riboviruses associated with microbial consortia dominated by bacteria and archaea in several acidic hot springs in Japan, to survey prokaryotic RNA virome. Whole genome sequences of two groups of riboviruses with multisegmented RNA genomes were obtained, one of which consists of unusual viruses with distinct RdRPs that seems to be intermediates between typical ribovirus RdRPs and RTs, whereas the other one is a distinct branch of partiti-like viruses. All genome segments in both these groups of viruses display the organization typical of bacterial riboviruses, where multiple genes encoding individual proteins are preceded by RBS.

## Materials and Methods

### Sample collection

A total of 11 samples were collected from five hot springs regions at southern Japan, in close proximity to active volcanoes (Table 1, Supplemental Information), according to the instructions of Unzen City, Unzen Nature Conservation Bureau and private companies that maintain each hot spring region. Temperature, pH and dissolved oxygen (DO) were measured in situ by using a multiple electrode sensor (D-55; Horiba, Kyoto, Japan). Concentration of H_2_S was determined by spectrophotometric approach of absorbance at a wavelength of 680 nm of methylene blue formed from the reaction with N,N-dimethyl-p-phenylenediamine in FeCl_2_-HCl solution. Typical measurement errors are 0.1 for pH, 0.1 mg/L for DO, and 5% for H_2_S. Dissolved chemicals and water isotope ratios of the geothermal waters were also measured and summarized in Supplemental Information.

**Table 1:**
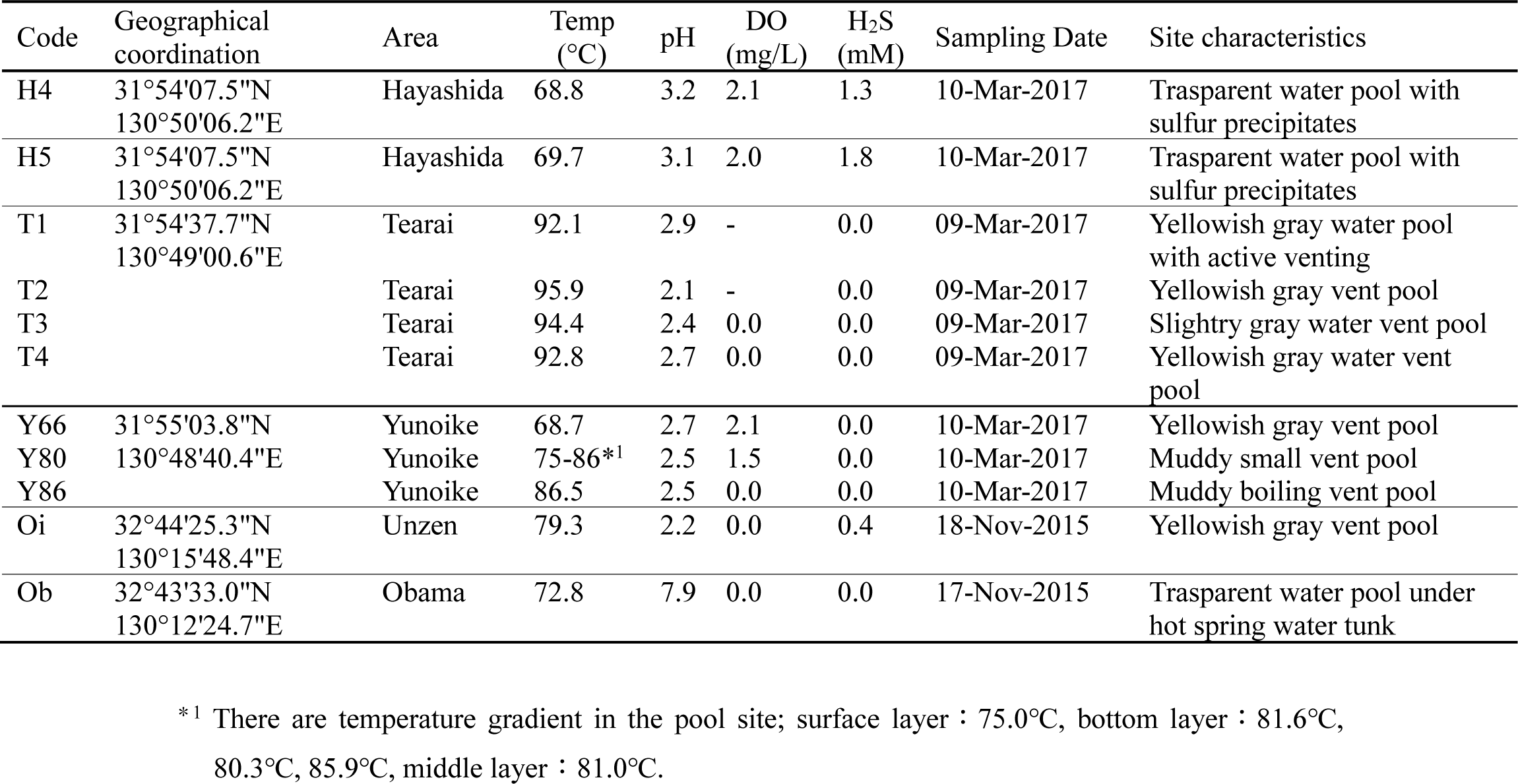
Characteristics of hot spring water samples.

Most of the sampling sites were characterized by high temperatures above 65°C, acidic (pH=2-3), and lower level of DO with accompanying by gray mud or light-yellow sulfur deposits. Site Ob was an only one exception with a slightly alkaline pH of pH=7.9. At each sampling station, approximately 10 L of hot spring water was collected in a sterilized plastic bag, and then filtered with 0.2-μm-pore-size cellulose acetate membrane filters in 47 mm diameter (Advantec, Tokyo, Japan) within 0.5-3 hours after sampling. The filters were stored at −80°C until nucleic acid extraction.

### RNA extraction

Cells collected on a portion of the 0.2-μm-pore-size filters corresponding to approximately 2 L of hot spring water were pulverized in a mortar in liquid nitrogen and suspended in dsRNA extraction buffer [20 mM Tris–HCl, pH 6.8, 200 mM NaCl, 2 mM EDTA, 1% SDS and 0.1% (v/v) β -mercaptoethanol] or TRIzol buffer for ds- and ssRNA purification, respectively. For dsRNA purification, total nucleic acids were manually extracted with SDS-phenol. dsRNA was purified using the cellulose resin chromatography method ^20,21^ as described previously ^19^. For ssRNA purification, the ssRNA fraction was collected using the TRIzol Plus RNA Purification Kit (Invitrogen, CA, USA) according to the manufacturer’s protocol. The ssRNA fraction was treated with DNase I (Invitrogen) and concentrated using the RNA Clean and Concentrator-5 Kit (Zymoresearch, CA, USA).

### cDNA synthesis

cDNA was synthesized from purified dsRNA and ssRNA as described previously ^19^. In brief, purified dsRNA was physically fragmented into about 1.5kbp and adapter oligonucleotide was ligated to 3’-end of fragmented dsRNAs. After heat denaturation with an oligonucleotide primer, which have complementary sequence to the adapter oligonucleotide, cDNA was synthesized using SMARTer RACE 5’/3’ Kit (Takara Bio, Shiga, Japan). ssRNA was converted into cDNA using SMARTer Universal Low Input RNA Kit according to the manufacturer’s protocol (Takara Bio). After PCR amplification, cDNA was fragmented by an ultrasonicator Covaris S220 (Covaris, MA, USA).

### Illumina sequencing library construction and sequencing

Illumina sequencing libraries were then constructed using KAPA Hyper Prep Kit Illumina platforms (Kapa Biosystems, MA, USA) from the physically shared environmental cDNAs. The libraries were sequenced using the Illumina MiSeq v3 Reagent Kit (600 cycles) with 300-bp paired-end reads on the Illumina MiSeq platform.

### Data processing

Cleanup reads were obtained using a custom Perl pipeline script (https://github.com/takakiy/FLDS) from dsRNA raw sequence reads ^17^. The clean reads were subjected to *de novo* assembly using CLC GENOMICS WORKBENCH version 11.0 (Qiagen Japan, Tokyo, Japan) with the following parameters: a minimum contig length of 500, word value set to auto, and bubble size set to auto. The full-length sequences were manually extracted using CLC GENOMICS WORKBENCH version 11.0 (Qiagen Japan), Genetyx version 14 (Genetyx, Tokyo, Japan), and Tablet viewer version 1.19.09.03 ^22^. as described previously ^23^. In this study, major full-length sequences with more than 1,000 average coverage were analyzed, except for Oi sample. For Oi sample, all full-length sequences were recovered. From ssRNA raw sequence reads, cleanup reads were also obtained using a custom Perl pipeline script (https://github.com/takakiy/FLDS). The resultant clean reads were applied to phyloFlash ^24^, to identify active microbes in our samples.

### Sequence analyses

RNA viral genes were identified using the BlastX program against the NCBI non-redundant (nr) database with an e-value ≤1×10^−5^. The ribosome-binding Shine-Dalgarno motifs were identified using Prodigal ^25^. Remote homology searches were performed using HHpred against the PDB70, Pfam, UniProt-SwissProt-viral70 and NCBI-CD (conserved domains) databases ^26^. Multiple sequence alignment of HsRV L_ORF4s was built using MEGA6. The alignment was then used as input in HHblits 3.3.0 which compared the alignments to the PDB70 (pdb70_from_mmcif_220313) database. Transmembrane domains were predicted using TMHMM ^27^.

### Modeling protein structures with AlphaFold2 and structural comparisons

Structural predictions for HsRV1 and HsRV-like RdRP amino acid sequences were performed using ColabFold 1.5.1 installed locally through LocalColabFold (https://github.com/YoshitakaMo/localcolabfold). A custom multiple sequence alignment with ten HsRV (HsRV1_La∼d, H5_contig_1, Oi_contig_1, Oi_contig_3, Oi_contig_5, Oi_contig_8, Oi_contig_9) and five HsRV-like (BDQA01000957, BDQA01004869, Ga0456180, Ga0393213, Ga0169446) RdRP amino acid sequences was used as an input. The number of recycles used for HsRV1_La ORF4 and HsRV-like RdRP predictions were 6 and 10, respectively. For the core (motifs A-C) region of marine HsRV-like RdRP BDQA01004869 (Fig. 3d), 20 recycles were used. The RdRPs of HsPV-H5 and pepper cryptic virus 1 (GenBank id: YP_009466859) were modeled using AlphaFold 2 through ColbFold v1.5.2 ^28,29^ with 6 recycles each. For the HsPV-H4 CP modeling, an alignment of RNA2 ORF1 homologs from HsPV-like viruses and Driatsky virus was used as a template with 12 recycles. The obtained model had a medium quality (average pLDDT=57.3), although the central region was modeled with higher quality (average pLDDT>70). This model was used as a query in DALI search which identified the CP of PCV1 (PDB id: 7ncr) as the best hit with the Z-score of 6.5. Thus, to improve the quality of the HsPV-H4 CP model, we repeated the modeling using the same sequence alignment and providing the PDB structure of the PCV1 CP as a template, with 24 recycles. The obtained model had an average pLDDT score of 78.1. Model display, structural alignment, coloring, and figure preparation were performed using UCSF ChimeraX software **^30^**.

### Phylogenetic analysis

Amino acid sequences of RdRP encoded by identified viruses and viruses related to the family *Partitiviridae* were aligned using MUSCLE ^31^ in MEGA6 ^32^. The ambiguous positions in the alignment were removed using trimAl with the option gt=0.8 ^33^. The RAxML program ^34^ was used for maximum likelihood-based phylogenetic analyses with a best-fit amino acid substitution model based on Akaike’s information criterion by ProtTest ^35^. Bootstrap tests were conducted with 1,000 samplings. To visualize phylogenetic trees, FigTree (version 1.4.2) and MEGA6 were used. For phylogenetic analysis of the HsRV-like RdRPs, the proteins were aligned using PROMALS3D ^36^, uninformative positions we removed using TrimAl with the *gappyout* functions ^33^. The final alignment contained 520 positions. The maximum likelihood tree was constructed using IQ-TREE v2 ^37^. The best-fitting substitution model was selected by IQ-TREE and was LG+I+G4. Branch supports were estimated using bootstrap (1,000 replicates).

### Data accessibility

Datasets obtained in this study have been available in the GenBank database repository (Accession Nos. HsRV1: BTCNxxxxxxxx-BTCNxxxxxxxx; HsPV-H4: BTCOxxxxxxxx-BTCOxxxxxxxx; HsPV-H5: BTCPxxxxxxxx-BTCPxxxxxxxx; HsPV-Y66: BTCQxxxxxxxx-BTCQxxxxxxxx; H5_contig_1: BTCRxxxxxxxx; Oi_contig_1-9: BTCSxxxxxxxx-BTCSxxxxxxxx) and Short Read Archive database (Accession No. DRAxxxxxx).

## Results and Discussions

### Composition of SSU rRNA and identification of RNA virus sequences associated with the hot spring microbiomes

To know the composition of active microbial consortia in the hot spring water samples, total ssRNA sequencing (ssRNA seq) reads were mapped on the small subunit (SSU) ribosomal RNA (rRNA) sequences from the Silva database (SILVA SSU version 138) using phyloFlash ^24^ (Fig. 1). Eukaryotic SSU rRNA reads represented less than 10% of the microbial communities in all samples. Especially for H4, H5, Y66 and Oi samples where RNA viruses were identified, eukaryotic SSU rRNA reads were less than 1% (Table S1). The microbial communities of T1, T2, T4 and Y86 samples, collected from the hot springs with temperatures above 85°C and pH<3, were predominated by archaea (>80% of total SSU rRNA reads), with thermoacidophilic archaeal family Sulfolobaceae representing >50% of the total SSU rRNA reads. By contrast, sequences related to thermoacidophilic bacterial family Hydrogenobaculaceae predominated (> 50 % of total SSU rRNA reads) in samples H4, H5, Y66 and Y80. The microbial community structures in the remaining three stations (T3, Oi and Ob) were distinct from those in the other samples. In particular, betaproteobacterial family Comamonadaceae occupied 25% of total SSU rRNA reads in Oi sample, which were negligible in other samples (Table S1).

**Fig. 1:**
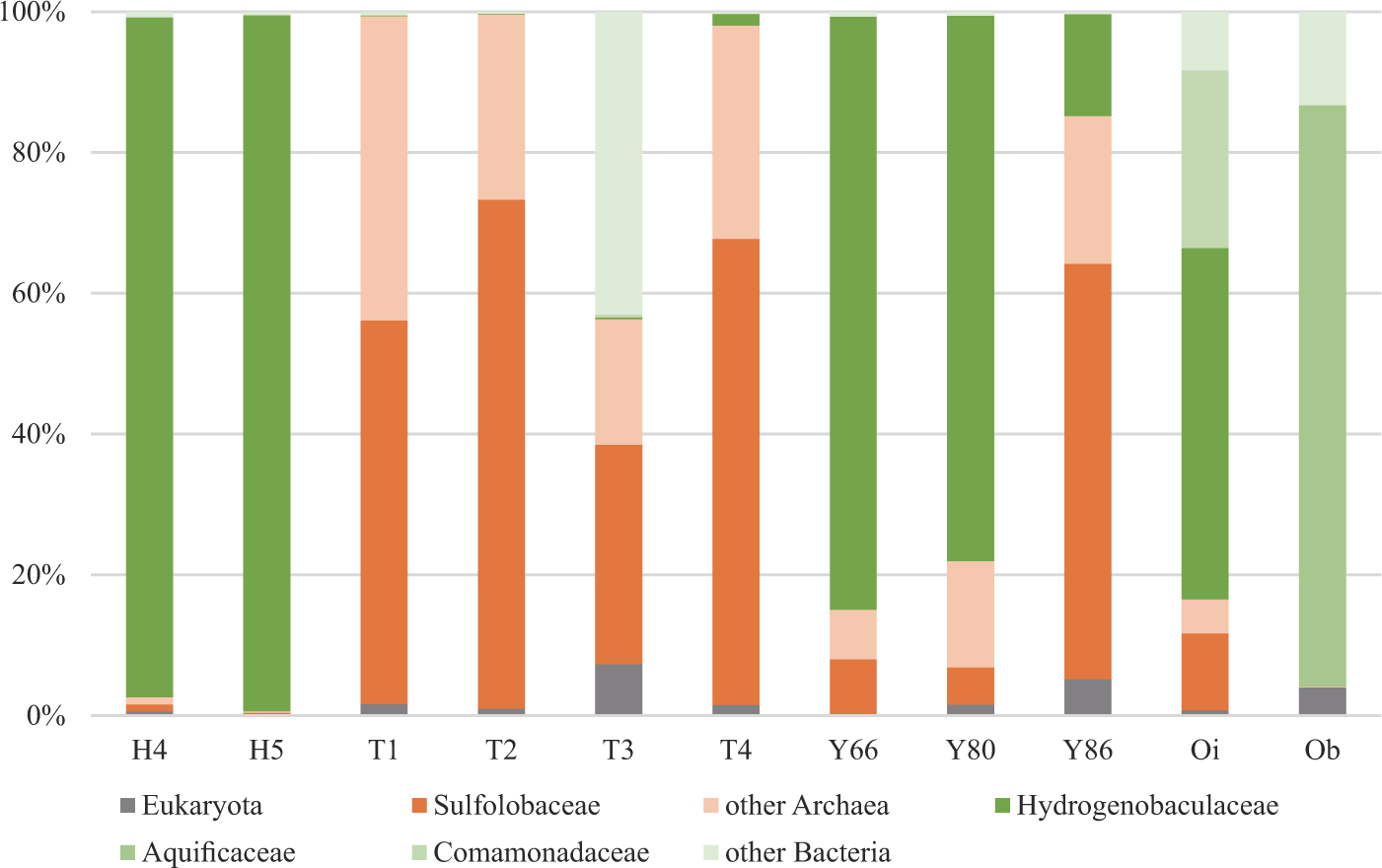
Composition of microbiomes associated with the hot spring samples. The composition was analyzed based on the mapped sequence reads on the rRNA sequences using phyloFlash. Details are given in Table S1.

In FLDS dsRNA sequencing (dsRNA seq), potential complete genomes of multipartite RNA viruses were obtained from samples H4, H5, Y66 and Oi (Table S2). For the samples from the other stations, successful sequence libraries were constructed except for Ob sample, but no contigs representing features of potential complete genomes of RNA viruses in FLDS read mapping ^18^ were obtained.

### Unusual tripartite RNA virus genomes from the Oi hot spring

The FLDS dsRNA seq of the Oi sample (79.3°C, pH 2.2) yielded three populations of contigs (Fig. 2a) which collectively recruited ∼50% of the clean FLDS reads from the Oi library. Among the contigs, we identified similar 5’ and 3’-terminal sequences (Fig. 2b), a feature characteristic of segmented RNA viruses, which facilitates the concerted encapsidation as well as replication of the multiple genomic segments (e.g., ^38^). Based on the similarity of the 5’ and 3’-terminal sequences, lengths of the segments, and gene content, we concluded that the three sets of contigs constitute genomes of a distinct group of tripartite RNA viruses. The segments were denoted L, M and S, for large, medium and small, respectively. Due to high sequence similarity between segment termini, we could not assign sets of segments to particular virus strains and thus analyzed them as a metapopulation. We obtained complete sequences for 4, 4 and 2 divergent variants of segments L, M and S, respectively (Fig. 2a).

**Fig. 2:**
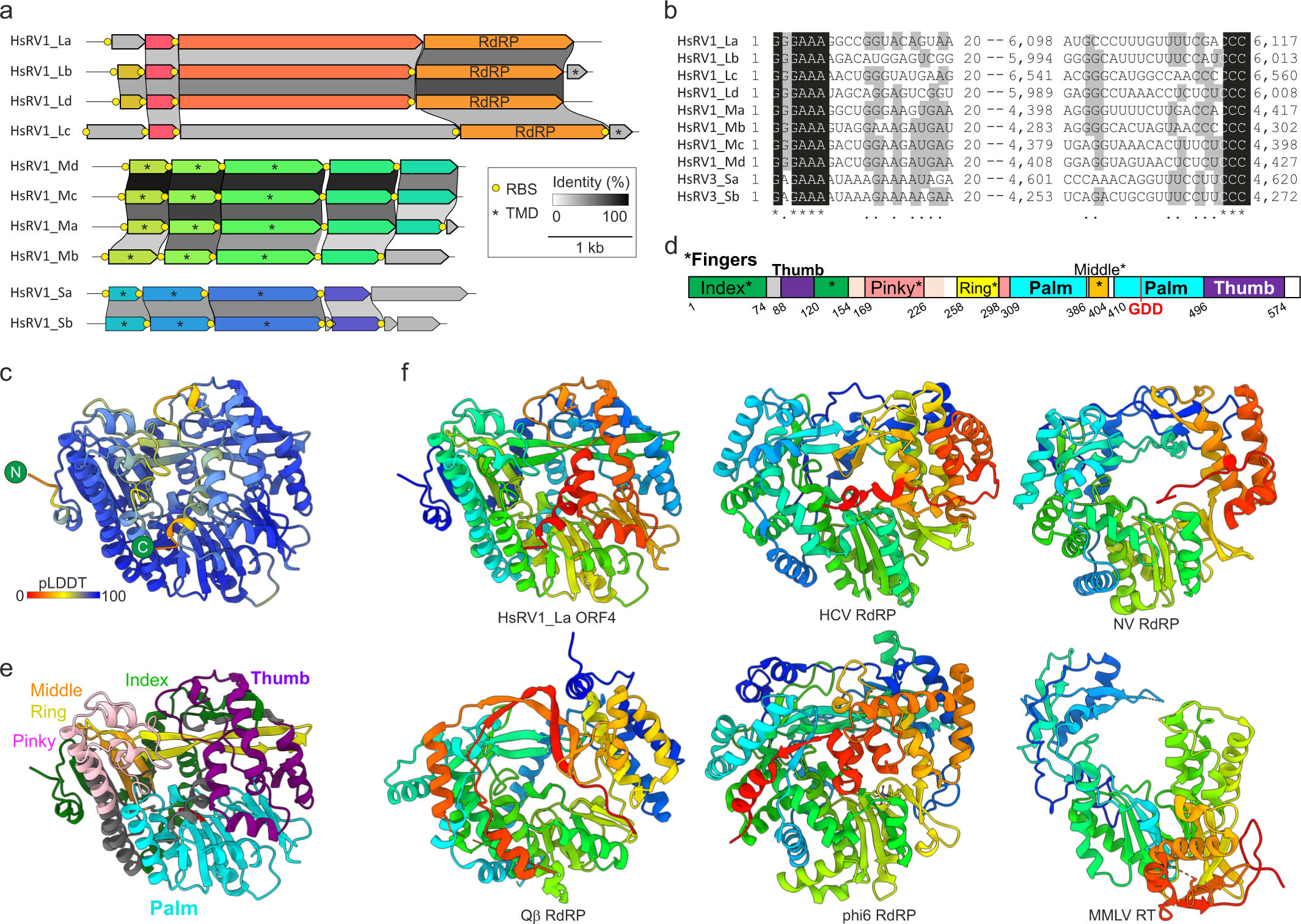
Unusual tripartite RNA virus genomes from the Oi hot spring. **a,** Genome organization and conservation of the three genomic segments (L, M and S) of HsRV1. Open reading frames encoding homologous proteins are shown as arrows with identical colors. Yellow circles represent predicted Shine-Dalgarno ribosome-binding sites (RBS). Asterisks denote putative genes encoding predicted transmembrane domain (TMD)-containing proteins. **b,** Multiple sequence alignment of the 5ʹ- and 3ʹ-terminal regions of the coding strands of reconstructed genome segments. Black shading, 100% nucleotide identity; gray shading, > 50% nucleotide identity. **c,** Quality assessment of the AlphaFold2 (AF2) model of the HsRV1 RdRP. The structural model is colored based on the per-residue Local Distance Difference Test (pLDDT) scores (average pLDDT=90.7), with the color key shown at the bottom left corner. **d-e,** Domain organization of the HsRV1 RdRP. **d,** Schematic representation of the domain organization, with exact coordinates of each subdomain, including the five ‘Fingers’, indicated. e, The structural model of HsRV RdRP colored using the same scheme as in panel d. **e,** Comparison of the HsRV RdRP with homologs from other RNA viruses, including hepatitis C virus (HCV; PDB: 6GP9), Norwalk virus (NV; PDB: 1SH0), Qbeta (PDB: 3MMP), phi6 (PDB: 1HHS) as well as reverse transcriptase (RT) from Moloney murine leukemia virus (MMLV; PDB: 4MH8). The structures are colored using the rainbow scheme from blue N-terminus to red C-terminus.

Segments L, M and S harbored 4-5, 5-6 and 5-7 open reading frames (ORFs), respectively (Fig. 2a). No ORFs were shared between segments of the three size classes, but within each class, the segments shared multiple ORFs (Fig. 2a; Table S3), indicating that the viruses from the Oi sample constitute an evolutionarily coherent lineage. Notably, M and S segments encoded three conserved predicted transmembrane proteins each (Fig. 2a), which could be involved in virus-host interactions^39^. None of the putative proteins encoded by the newly discovered viruses showed similarity (E-value=5e-03) to any protein sequences in public databases. Even most sensitive profile-profile searches using HHpred yielded no significant (HHpred probability >90%) hits for any of the predicted proteins. However, HHpred searches queried with the amino acid sequence of ORF4 from the L segment produced a partial hit to RdRPs of RNA viruses. Although the hits were not significant (HHpred probability <90%) and encompassed only a small region of the RdRP (∼15% of the target profile), the aligned region covered the diagnostic RdRP motifs B (SGxxxT, x – any amino acid) and C (GDD) (Fig. S1), and we thus decided to further pursue this clue.

HMMscan ^40^ using Neo-HMM (score >20) ^41^ and RVDB-HMM (score >20) ^42^, and palmscan ^43^ did not identify L_ORF4 as an RdRP. Furthermore, no sequence related to L_ORF4 could be identified using PZLAST ^44^ among several tera-bytes of public metagenomic sequences either. Similarly, attempts to generate a structural model of L_ORF4 using AlphaFold2 with a default multiple sequence alignment (MSA) generation method did not result in a reliable model, likely due to the lack of identifiable homologs in public databases. Thus, we set out to enrich the sequence diversity of L_ORF4 by reanalyzing the all FLDS data. To this end, the unmapped sequence reads were assembled and L_ORF4 protein sequences were used as queries to search against the assembled contigs using BLASTX. This search yielded 10 additional L_ORF4-like sequences encoded by H5_contig_1 from H5 and Oi_contigs_1-9 from Oi samples (E-value ≤ 1 × 10^−5^) (Table S4). The additional homologs detected in this search were combined with the 4 initially identified L_ORF4 sequences and the produced MSA was used as a query in HHpred search against the PDB70 database. This search yielded significant hits (probability >90%) to RdRPs from various RNA viruses, although the aligned region remained limited (∼15% of the target profiles). Collectively, these searches suggested that L_ORF4 homologs are highly divergent RdRPs.

Using the MSA that included the identified L_ORF4 homologs, a high quality (average pLDDT=90.7) AF2 model of the putative RdRP was obtained (Fig. 2c). Examination of this model revealed a topology typical of the palm-domain polymerases, with readily discernible ‘Fingers’, ‘Palm’ and ‘Thumb’ subdomains (Fig. 2d, e) and the overall architecture similar to that of viral RdRPs (Fig. 2f), albeit with some unique structural features. In particular, L_ORF4 model displayed an extended and highly ordered ‘Fingers’ subdomain, with the ‘fingertips’ forming a 5-stranded β-sheet not present in other RdRPs which interacts with the ‘Thumb’ subdomain. The conserved motifs B and C identified using the HHpred analysis were located within the Palm subdomain, at the position equivalent to that in other RdRPs. Furthermore, structural superposition of the Palm subdomains from different RdRPs allowed identification of the third core motif, A, in L_ORF4 (see below). Thus, we concluded that L_ORF4 represents an RdRP and accordingly provisionally named the discovered tripartite virus “Hot spring RNA virus (HsRV)”. The four RdRPs encoded by the complete L segments display only 37 to 75% pairwise identity and thus can be considered to represent four distinct virus species (or even higher taxa).

### HsRV-like viruses are also present in non-extreme (moderate) aquatic ecosystems

The obtained sequence profile of the HsRV RdRP was used to search the previously described FLDS sequence data from the coastal seawater samples ^19^, which led to the identification of two additional HsRV-like RdRP-encoding contigs (GenBank accessions: BDQA01000957 and BDQA01004869). The two incomplete RdRP sequences were encoded on short contigs (1.2 and 1.4 kb). Nevertheless, searches against the IMG/VR database queried with the corresponding RdRPs yielded significant hits (E-value ≤ 1 × 10^−5^) to three additional RdRP sequences encoded on apparently complete or near-complete 5.3-5.6 kb-long genome segments (Ga0456180_000042, Ga0393213_00017, Ga0169446_00510; Fig. 3a, S2 and Table S6).

**Fig. 3:**
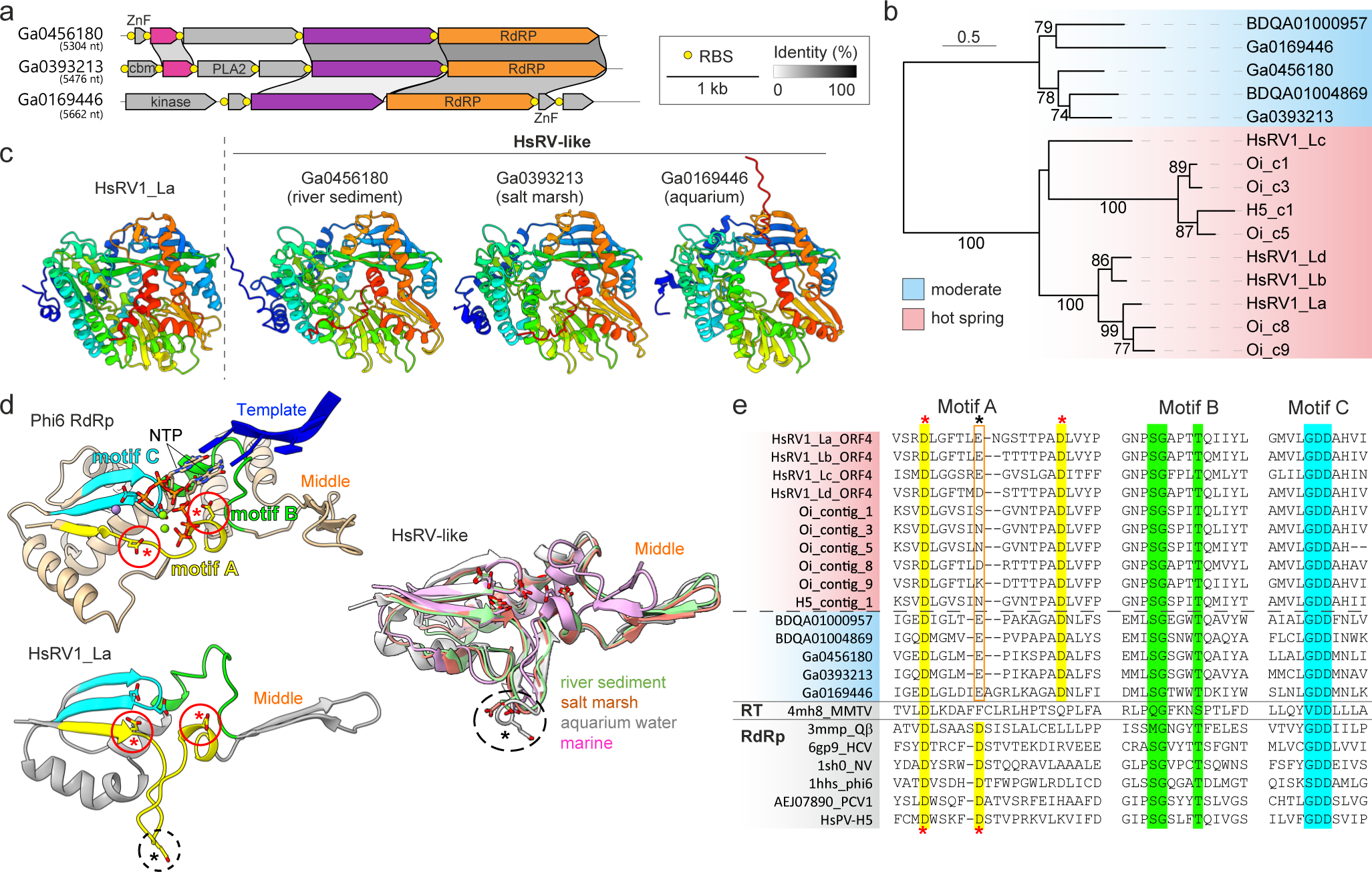
HsRV-like viruses from moderate environments. **a,** RdRP-encoding segments of HsRV-like viruses from non-extreme aquatic ecosystems. Open reading frames encoding homologous proteins are shown as arrows with identical colors. Yellow circles represent predicted Shine-Dalgarno ribosome-binding sites (RBS). Asterisks denote putative genes encoding predicted transmembrane proteins. **b,** Maximum likelihood phylogeny of the HsRV-like RdRPs encoded by viruses from extreme (pink) and moderate (blue) ecosystems. Branch support was assessed using the approximate likelihood-ratio test (aLRT), with the corresponding values (%) shown on the branches. The scale bar represents the number of substitutions per site. **c,** Comparison of the HsRV1 RdRP with the homologs encoded by viruses from moderate aquatic ecosystems. The models are colored using the rainbow scheme from blue N-terminus to red C-terminus. **d,** Comparison of the catalytic cores encompassing the conserved RdRP motifs A (yellow), B (green) and C (cyan). Top: the structure of bacteriophage phi6 RdRP with the substrate nucleoside triphosphates (NTP) and template RNA strand (blue ribbon). Bottom: the HsRV1 RdRP. Middle: structurally superposed HsRV-like RdRPs from moderate ecosystems. The NTP and active site residues of motifs A and C are shown using the stick representation. The conserved aspartate residues of Motif A are circled, with structurally equivalent residues indicated with red asterisks, whereas the non-conserved residue located in the loop facing away from the Motif C in HsRV1 and related RdRP is indicated with the black asterisk. **e,** Multiple sequence alignment of the conserved Motifs A, B and C of HsRV-like RdRPs from extreme (red shading) and moderate (blue shading) ecosystems with the corresponding regions from RdRPs and reverse transcriptase from other viruses (grey shading), including Moloney murine leukemia virus (MMLV), hepatitis C virus (HCV), Norwalk virus (NV), Pepper cryptic virus 1 (PCV1) and hot spring partiti-like virus H5 (HsPV-H5). The sequences are indicated with the PDB or GenBank accession numbers. The conserved residues in Motifs A, B and C are shaded yellow, green and cyan, respectively, matching those in panel d. The conserved aspartate residues of Motif A are highlighted in yellow, with structurally equivalent residues indicated with red asterisks, whereas the non-conserved residue in HsRV-like RdRPs located at the equivalent position as the second aspartate in other RdRPs is indicated with the black asterisk.

Similar to HsRV, the RdRPs of these related viruses from moderate aquatic environments did not show similarity to sequences in current databases either using BLASTP or HHpred searches. However, the functions of several other proteins could be predicted from HHpred search results. In particular, ORF1 and ORF3 of Ga0393213 encode putative carbohydrate-binding protein with the jelly-roll fold and a phospholipase A2 (PLA2), respectively. Notably, PLA2 has been previously identified in viruses with small RNA and ssDNA genomes ^4,45^ and, in parvoviruses, shown to be important for infectivity ^45^. Ga0169446 encodes a predicted kinase related to nucleoside monophosphate kinases. Finally, Ga0169446 and Ga0456180 encode non-orthologous zinc finger proteins, which could be involved in nucleic acid binding or protein-protein interactions.

Apart from the RdRP, the HsRV-like viruses lack predicted proteins shared with HsRVs from hot springs. However, the three viruses from the moderate environments share a ∼500 aa-long protein encoded upstream of the RdRP, a likely candidate for the capsid protein (Fig. 3a). In addition, Ga0456180 and Ga0393213 share a protein of unknown function.

Ga0456180, Ga0393213 and Ga0169446 originate from floodplain (river sediments), salt marsh, and aquarium samples, respectively. Maximum likelihood phylogenetic analysis of HsRV-like RdRPs showed clear separation between viruses from the hot spring and those from moderate aquatic environments (Fig. 3b). Collectively, these results suggest that HsRV-like viruses are broadly distributed in aquatic ecosystems.

### Structural analysis confirms the uniqueness of the HsRV-like RdRPs

To further analyze the features of HsRV-like RdRPs and their relationship to RdRPs encoded by other viruses, we generated AF2 models for the three HsRV-like RdRPs from moderate ecosystems (Fig. 3c). The obtained models showed overall structural similarity with the RdRP of HsRV1, including the extended ‘Fingers’ subdomain (Fig. 3c). Further analysis of the multiple sequence and structure alignments revealed another unusual feature of these proteins, namely, a unique, extended RdRP motif A. In the canonical motif A, the two conserved Asp residues involved in catalysis and substrate discrimination ^46,47^, respectively, are separated by 4-5 residues and bracket the catalytic GDD residues of motif C (Fig. 3d, e). By contrast, in HsRV-like RdRPs, the second Asp residue of motif A is not conserved, and the corresponding residue is located in a loop facing perpendicularly away from motif C, suggesting that it cannot perform the same function. However, all analyzed HsRV-like RdRPs contain an Asp (Asp*) which is located 12-14 residues away from the first Asp of motif A (Fig 3e). Analysis of the structure showed that, despite the extended spacing in the protein sequence, Asp* occupies a position equivalent to the second Asp of the canonical motif A (Fig. 3d, e) and is likely to be its counterpart involved in substrate discrimination. Such extended motif A has not been reported in other RdRPs and is one of the defining features of the HsRV-like RdRPs.

We next performed structural clustering based on the pairwise DALI Z-scores of the HsRV-like RdRPs together with RdRPs of a selection of other RNA viruses and structurally-related reverse-transcriptases (RT) encoded by eukaryotic viruses of the order *Ortervirales* ^48^ as well as non-viral RTs from bacteria and eukaryotes (Fig 4a). In this analysis, HsRV-like RdRPs from both hot springs and moderate aquatic ecosystems formed a tight cluster, underscoring their relatedness despite high sequence divergence. Notably, whereas all previously known viral RdRPs formed a clade in the structure-based dendrogram, the HsRV-like RdRPs remained separated from those (Fig. 4a). In particular, the two viral RdRP clusters were interspersed with the RTs, such that the viral RTs were the closest structural neighbors of the HsRV-like RdRPs. This result further confirms the extreme divergence of the HsRV-like RdRPs and might reflect a closer relationship to viral RTs. This unexpected link was further strengthened by the comparison of the ‘Palm’ subdomain of HsRV-like RdRPs with homologs from other RNA viruses as well as viral and non-viral RTs. In RdRPs of RNA viruses from 5 established phyla ^10^, the first β-strand (blue in Fig. 4b) containing motif A and the motif B-containing α-helix are separated by a characteristic helix-turn-helix (HTH) region followed by a β-hairpin corresponding to “Middle” fingers subdomain (Fig. 2d, e). However, the HTH motif is absent in both the HsRV-like RdRPs and viral RTs. Notably, non-viral RTs, such as those from group II introns or retrons, contain the HTH motif but lack the β-hairpin region, which is compatible with the intermediate position of RTs between the two clades of viral RdRPs. Furthermore, the HsRV-like RdRPs might comprise evolutionary intermediates between viral RTs and RdRPs.

**Fig. 4:**
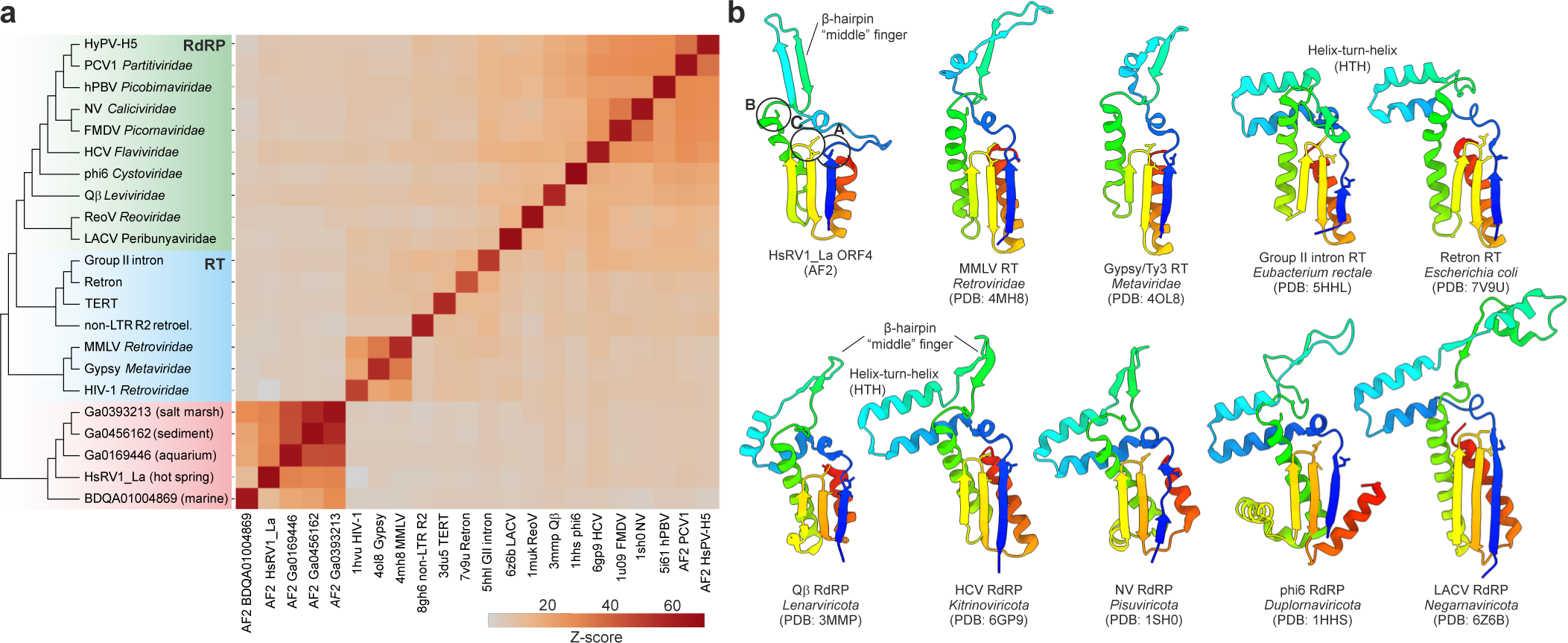
Structural relationships between RdRPs and RTs. **a,** Matrix and cluster dendrogram were constructed based on the pairwise Z-score comparisons calculated using DALI. Different protein groups are highlighted with different background colors on the dendrogram: RdRPs from previously characterized viruses, green; viral and non-viral reverse transcriptases (RT), blue; HsRV-like RdRPs, red. The color scale indicates the corresponding Z-scores. Moloney murine leukemia virus (MMLV), hepatitis C virus (HCV), Norwalk virus (NV), Pepper cryptic virus 1 (PCV1) and hot spring partiti-like virus H5 (HsPV-H5), hPBV, human picobirnavirus; FMDV, foot-and-mouth disease virus; ReoV, reovirus; LACV, La Crosse virus; HIV-1, human immunodeficiency virus 1; TERT, telomerase reverse transcriptase; non-LTR R2 retroel., non-long terminal repeat R2 retroelement; AF2, AlphaFold2 model. For experimentally determined structures, the corresponding PDB accession numbers are indicated at the bottom of the matrix. **b,** Structural comparison of the core domain of RdRPs and RT encompassing the conserved motifs A-C. The structures are colored using the rainbow scheme from blue N-terminus to red C-terminus.

### A thermoacidophilic partiti-like virus

Analysis of the FLDS RNA sequencing data from the stations H4 (68.8°C, pH 3.2), H5 (69.7°C, pH 3.1) and Y66 (68.7°C, pH 2.7) revealed a bipartite virus genome unrelated to HsRV1 (Fig. 5a). The virus contigs recruited 82%, 95% and 12% of non-rRNA trimmed FLDS reads from the H4, H5 and Y66 libraries, respectively (Table S2). Although Y80 data also contained related sequences, they were incomplete. The genomic segments, RNA1 and RNA2, shared conserved 5’ terminal sequences, and encoded one and two proteins, respectively (Fig. 5b). ORF1 of RNA was unambiguously identified as an RdRP, yielding significant BLASTP hits to RdRPs of members of the *Partitiviridae* family, with the best hit being to the unclassified Driatsky virus (QIS87951; E-value=1e-95). Accordingly, we tentatively named the virus hot spring partiti-like virus 1 (HsPV1). Due to high sequence similarity between segment termini, we could not assign sets of segments to particular virus strains and thus analyzed them as a metapopulation. Maximum likelihood phylogenetic analysis of the RdRP sequence from diverse partiti-like viruses showed that HsPV1 and Driatsky virus (see below) form a deeply branching clade which is a sister group to viruses classified into the genus *Deltapartitivirus* (Fig. 5c). AF2 modeling yielded an HsPV1 RdRP model closely similar to that of the RdRP of the deltapartitivirus Pepper cryptic virus 1 (Fig. 5d, Fig. S3a), which was confirmed by DALI Z-score-based clustering (Fig. 4a), where the two viruses formed a clade next to picobirnaviruses.

**Fig. 5:**
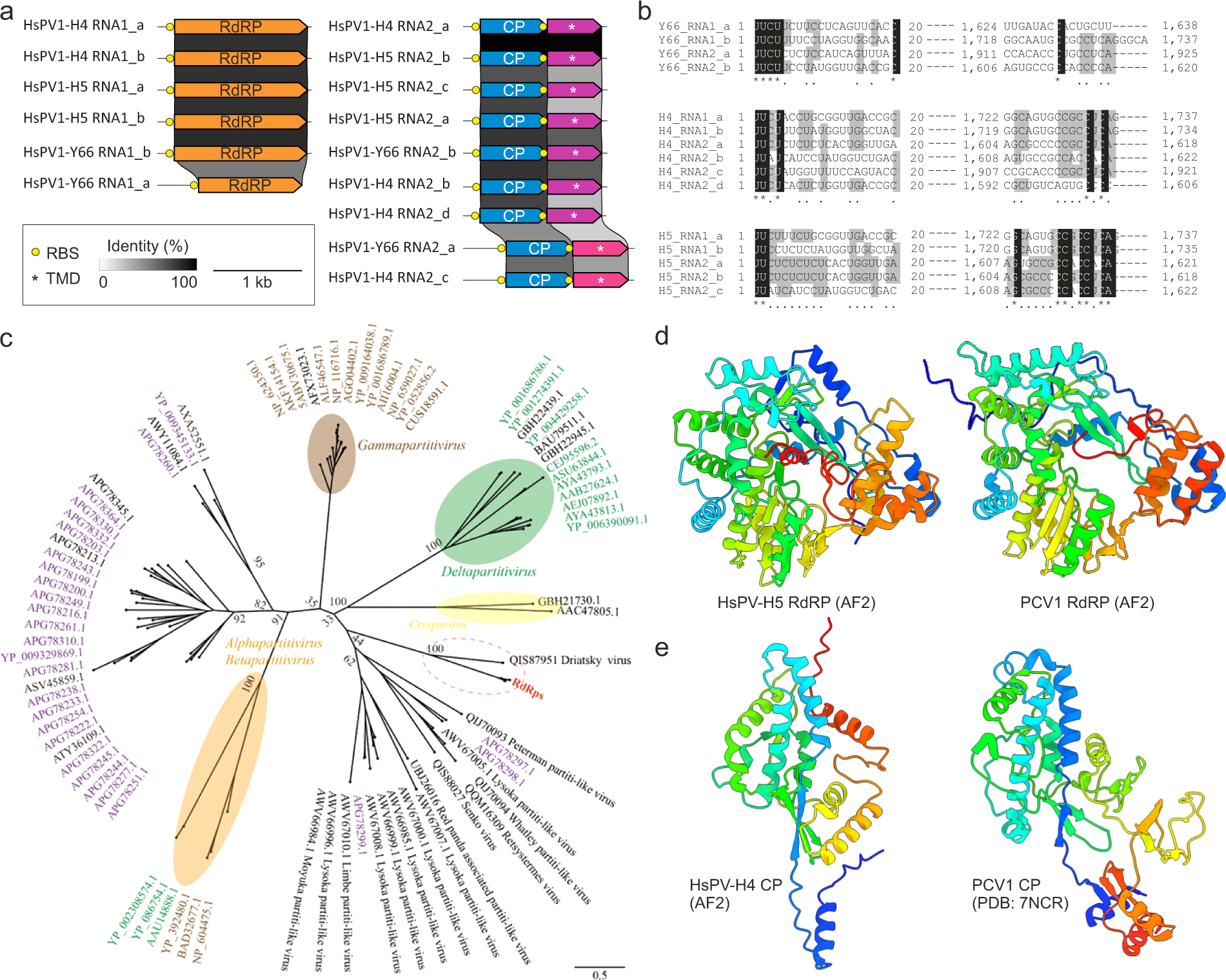
A thermoacidophilic partiti-like virus. **a,** Genome organization and conservation of the two genome segments, RNA1 and RNA2, of HsPV1. Open reading frames encoding homologous proteins are shown as arrows with identical colors. Yellow circles represent predicted Shine-Dalgarno ribosome-binding sites (RBS). Asterisks denote putative genes encoding predicted transmembrane domain (TMD)-containing proteins. RdRP, RNA-dependent, RNA polymerase; CP, capsid protein. **b,** Multiple sequence alignment of the 5ʹ- and 3ʹ-terminal regions of the coding strands of reconstructed genome segments. Black shading, 100% nucleotide identity; gray shading, > 50% nucleotide identity. **c,** Maximum likelihood phylogeny of the RdRP proteins from representative members of the family *Partitiviridae* and related sequences (including all HsPV strains). Sequences are indicated with the corresponding GenBank accession numbers. Numbers at the branches indicate the bootstrap support (%) from 1,000 RAxML bootstrap replicates. We used RAxML with the LG+G+I+F model. Colors of virus names indicate the classification of the host organism: green, plant; brown, fungi; purple, from ^2^; black, others or unclear. d, Comparison of the RdRP from HsPV-H5 with a homolog from deltapartitivirus Pepper cryptic virus 1 (PCV1). e, Comparison of the CP from HsPV-H4 with a homolog from deltapartitivirus PCV1. AF2, AlphaFold2 model. The structures are colored using the rainbow scheme from blue N-terminus to red C-terminus. The HsPV RdRP and CP structures colored based on the pLDDT quality scores can be found in Figure S2.

ORF1 and ORF2 of RNA2 did not show significant sequence similarity to proteins in sequence or profile databases, except that ORF2 was predicted to encode a membrane protein with two transmembrane domains. Given the relatively high similarity between the RdRP of HsPV1 and Driatsky virus (Accession: MT025082), we explored the possibility that the second genomic segment of Driatsky virus was missed, not an uncommon situation with segmented RNA viruses, especially for segments not encoding the RdRP. Driatsky virus was identified in a metatranscriptome of an environmental animal sample, but labeled to be “unlikely vertebrate associated”. To identify the potential RNA2 of Driatsky virus, we searched contig sequences from the corresponding metatranscriptome (SRR7239362) using HsPV RNA2 as a query. BLASTX analysis against ORF1 of RNA2 identified only one significantly similar sequence, contig_9778 (Supplemental Information). This contig encompassed two ORFs, as in the case of HsPV1 RNA2, and the amino acid sequences of contig_9778 ORF1 showed significant similarity (E-value = 2e-06) with HsPV-H4 RNA2_c_ORF1 protein. The phylogenetic relationship between those sequences is shown in Fig. S4.

Structural modeling of RNA2 ORF1 using a custom alignment that included orthologs from different HsPV1 strains and Driatsky virus yielded a high-quality model (pLDDT=78.8), with only the terminal regions being of lower quality (Fig. S3b). Structure-based searches against the PDB database using DALI produced significant hits to structurally related capsid proteins of partitiviruses and picobirnaviruses ^49-51^, with the best match (Z-score=8.2) to the capsid protein of Pepper cryptic virus 1 (Fig. 5e; PDB id: 7ncr; *Deltapartitivirus*). Thus, based on the phylogeny of the RdRP and structural similarity of the capsid protein, HsPV1 appears to be a genuine member of the family *Partitiviridae*. However, unlike in other classified members of the family, RNA2 of HsPV1 is bicistronic and encodes a membrane protein of unknown function, which is likely to participate in virus-host interaction.

### HsRV1 and HsPV1 likely infect prokaryotic hosts

All samples in which HsRV1 and HsPV1 were detected nearly exclusively contained RNA sequences from prokaryotes, with eukaryotic presence being below 1%. This is consistent with eukaryotes being unable to thrive in polyextremophilic conditions combining high temperatures and acidic pH. Notably, the microbial communities in all 4 samples (H4, H5, Y66 and Oi) were dominated by bacteria. Thus, it appears most likely that HsRV1 and HsPV1 infect bacteria. To test this inference, we predicted ribosome-binding Shine-Dalgarno (SD) motifs in all HsRV1 and HsPV1 strains using Prodigal. SD motifs are essential for translation initiation in many prokaryotes, and their conservation is a diagnostic feature of prokaryotic genes that has been used to assign bacterial hosts to several groups of RNA viruses, including picobirnaviruses and partitiviruses ^8,52^. Analysis of the HsRV1 and HsPV1 genomes showed that nearly every gene in these viruses is preceded by an SD motif (Fig. 2a, 3a, 5a, Table S5), further suggesting that both HsRV1 and HsPV1 infect prokaryotic hosts. Bacteria of the genus *Hydrogenobaculum* (family Aquificaceae) were predominant (>95%) in samples H4 and H5 and highly abundant in Y66 (>85%), suggesting that HsPV1 detected in all three samples infects *Hydrogenobaculum* sp.

It has been shown that RT-encoding CRISPR-Cas systems can acquire spacers from RNA virus genomes, thereby providing information on the host association of such CRISPR-targeted RNA viruses ^8^. No spacers matching the HsRV1 and HsPV1 genomes were identified in the public databases. Metagenomic DNA sequencing of the hot spring samples yielded 919 CRISPR spacer sequences, but none of them showed significant similarity to the identified RNA virus genomes either (Supplemental Information). Nevertheless, the lack of eukaryotes in the hot spring samples, contrasted by the dominance of bacteria, combined with the presence of typical prokaryotic SD motifs upstream of the predicted virus genes and the overall polycistronic organization of the viral genomes, collectively strongly suggest that HsRV1 and HsPV1 are viruses of thermophilic bacteria.

### Expansion of prokaryotic RNA virus diversity and habitat

This work continues the trend established by recent large scale metatranscriptome surveys that substantially expanded the prokaryotic RNA virome as well as the ecological range of RNA viruses ^7,8^. Unlike those large-scale studies, however, here we report complete genomes of multi-segmented riboviruses, adding information on their genome organization and gene content, beyond the RdRP. Some groups of riboviruses, such as picobirnaviruses and certain branches of partitiviruses, previously thought to infect eukaryotes, have been reassigned to prokaryotic hosts through a combination of evidence including their recovery from habitats that are heavily dominated by prokaryotes, genome organization, the presence of SD motifs, and in some cases, encoding of cell wall lysis enzymes and CRISPR targeting ^8^. Here we identify another group of partiti-like viruses that apparently infect a bacterial host, most likely, the extremely thermophilic *Hydrogenobaculum* sp. Notably, these HsPV1-like viruses appear to have a specific evolutionary affinity with Deltapartitiviruses that have well established eukaryotic hosts. Thus, this finding supports the previous conclusions on multiple switches from eukaryotic to prokaryotic hosts (and vice versa) among partiti-like viruses, and further, suggests that these viruses can also switch between mesophilic and thermoacidophilic hosts.

The discovery of the HsRV1-like group of riboviruses also recapitulates previous finding of several small groups of riboviruses that are predicted to infect bacteria and might become distinct phyla ^7,8^, given that they do not cluster with any of the established phyla in the phylogenetic tree of the RdRP. However, the RdRP of HsRV1 and its relatives seems to deviate from the RdRP consensus farther than any of the other putative phyla and possess unusual (predicted) structural features that appear to link them to viral RTs. Whether this connection reflects an intermediate position of the HsRV1-like viruses between the kingdoms *Orthornavirae* and *Pararnavirae*, or results from convergent evolution, remains uncertain and should be clarified by sequencing and structural analysis of additional members of this group, or possibly, other groups of riboviruses with similar features. Regardless, the HsRV1-like viruses are a strong candidate for a separate phylum in the kingdom *Orthornavirae,* or potentially, even a third kingdom within the realm *Riboviria*. Apart from the unusual RdRP, the distinctness of this group is emphasized by the lack of functional prediction for the rest of their proteins. In particular, there is no clarity as to the identity of their capsid proteins and accordingly, the virion structure that remains to be studied experimentally. We note, however, that ORF3 of segment L, which encodes the largest protein of the virus, is a likely candidate for one of the major capsid proteins. Similar to the inner capsid protein P1 of cystoviruses ^53^, the protein is encoded on the same segment (L) as the RdRP. The other conserved non-membrane proteins encoded on segments M and S could encode additional virion components. Similarly, the HsRV1-like viruses from moderate ecosystems also encode a conserved protein at the equivalent position, which could represent their capsid protein. Although the two groups of potential capsid proteins do not share appreciable sequence similarity and substantially differ in length (∼1000 aa vs ∼500 aa), we cannot exclude the possibility that they are extremely divergent homologs. Regardless, the divergence between the HsRV-like viruses from moderate and extreme environments would justify their classification into separate families unified within a higher rank taxon through the shared RdRPs.

The discovery of RNA viruses in acidic and high temperature conditions seems unexpected given the fragility of long RNA molecules. However, genome segmentation, with relatively small individual segments, and the likely double-stranded genome structure, which is associated with higher thermal stability ^54^, apparently obviate this problem. Given these findings, it becomes particularly puzzling why no RNA viruses have been so far linked to archaea. A ribovirus genome has been previously sequenced from a Yellowstone hot spring and tentatively assigned a hyperthermophilic archaeal host ^55^. Subsequently, however, numerous related ribovirus genomes have been identified in a mesophilic environment ^4^, whereas there was no follow-up on putative archaeal virus; thus, the status of this finding remains uncertain. One possibility is that such viruses remain unrecognized because they encode extremely diverged RdRPs or even distinct RdRPs unrelated to those encoded by the members of the *Riboviria*.

Taken together, this report is a proof of concept for the discovery of multiple, perhaps many groups of RNA viruses with unexpected properties by obtaining complete genomes of segmented RNA viruses from meta-dsRNA seq data and mining metatranscriptomes from habitats with distinct conditions. Information on non-RdRP segments is not available for most of the RNA virus lineages identified only from the metatranscriptome, and we cannot even recognize the existence of RNA viruses that are distantly related to known RNA viruses. Our concept overcomes these limitations for segmented RNA viruses and this will greatly contribute to making RNA virome information more complete.

## Acknowledgements

The authors thank Kirishima Iwasaki Hotel, NIPPON PAPER LUMBER CO., LTD. and NITTETSU MINING CO., LTD KAGOSHIMA GEOTHERMAL FACILITY for their support for field sampling. We thank Shinsuke Kawagucci, Mitsuhiro Yoshida, Yukari Yoshida-Takashima, Miwako Tsuda and Fumie Kondo for discussions, suggestions, sample collections and preliminary experiments related to this study. This study was supported by JSPS KAKENHI (Grant Nos. 15H05468, and 20K20377) and by Grants-in-Aid for Scientific Research on Innovative Areas from the Ministry of Education, Culture, Science, Sports, and Technology (MEXT) of Japan (Grant Nos. 22H04879, 20H05579, 19H05684, 16H06429, 16K21723, and 16H06437). This research was also supported in part by Lilly Endowment, Inc., through its support for the Indiana University Pervasive Technology Institute, and by a grant from the Institute for Fermentation, Osaka, Japan. E.V.K. is supported by the Intramural Research Program of the National Institutes of Health of the USA (National Library of Medicine).

---

**Table S1.** Relative abundances of rRNA reads in ssRNA seq of representative microbial lineages.

|  | H4 | H5 | T1 | T2 | T3 | T4 | Y66 | Y80 | Y86 | Oi | Ob |
| --- | --- | --- | --- | --- | --- | --- | --- | --- | --- | --- | --- |
| Eukaryota | 0.59 | 0.19 | 1.64 | 0.99 | 7.31 | 1.51 | 0.11 | 1.53 | 5.16 | 0.75 | 4.02 |
| Sulfolobaceae | 1.00 | 0.23 | 54.50 | 72.32 | 31.19 | 66.31 | 7.89 | 5.28 | 59.04 | 10.94 | 0.00 |
| other Archaea | 1.01 | 0.19 | 43.23 | 26.29 | 17.83 | 30.36 | 7.06 | 15.11 | 20.99 | 4.83 | 0.14 |
| Hydrogenobaculaceae | 96.57 | 98.90 | 0.13 | 0.12 | 0.27 | 1.61 | 84.24 | 77.54 | 14.47 | 49.84 | 0.01 |
| Aquificaceae | 0.00 | 0.01 | 0.00 | 0.00 | 0.00 | 0.00 | 0.01 | 0.00 | 0.00 | 0.09 | 82.54 |
| Comamonadaceae | 0.05 | 0.00 | 0.00 | 0.00 | 0.38 | 0.00 | 0.01 | 0.00 | 0.00 | 25.21 | 0.00 |
| other Bacteria | 0.78 | 0.47 | 0.50 | 0.29 | 43.03 | 0.32 | 0.69 | 0.56 | 0.34 | 8.32 | 13.29 |

**Table S2:**
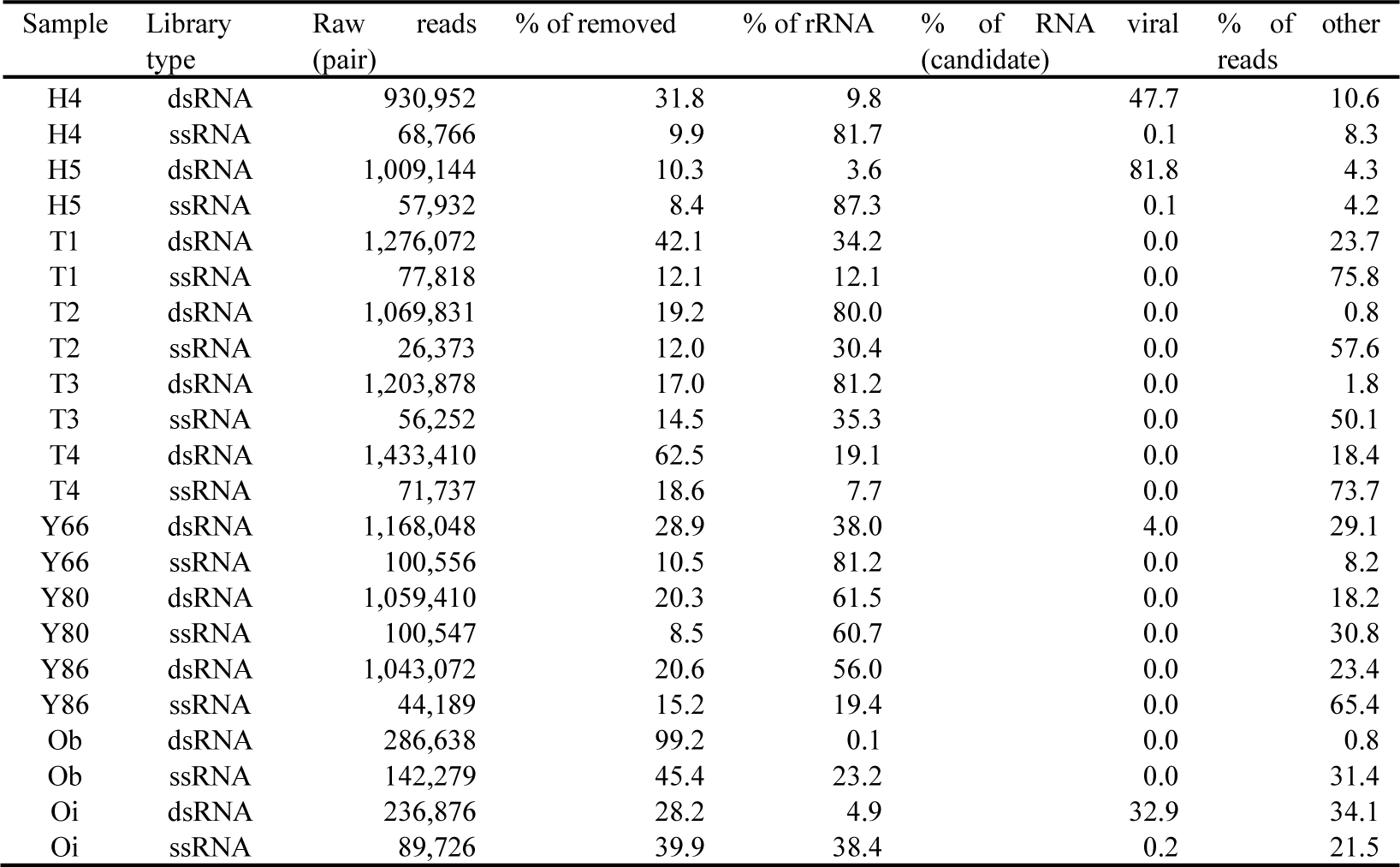
Classification of NGS reads.

**Table S3:**
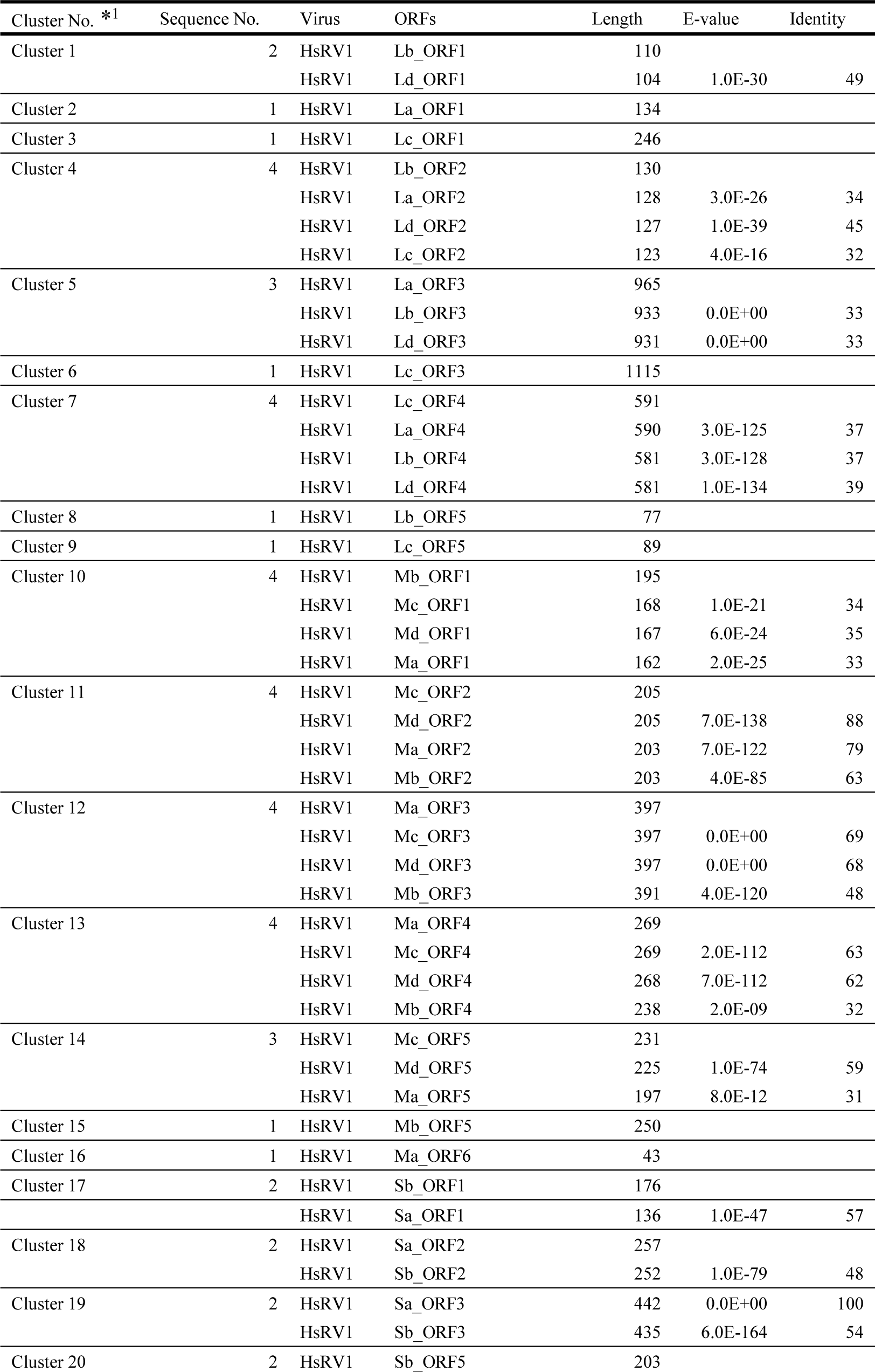
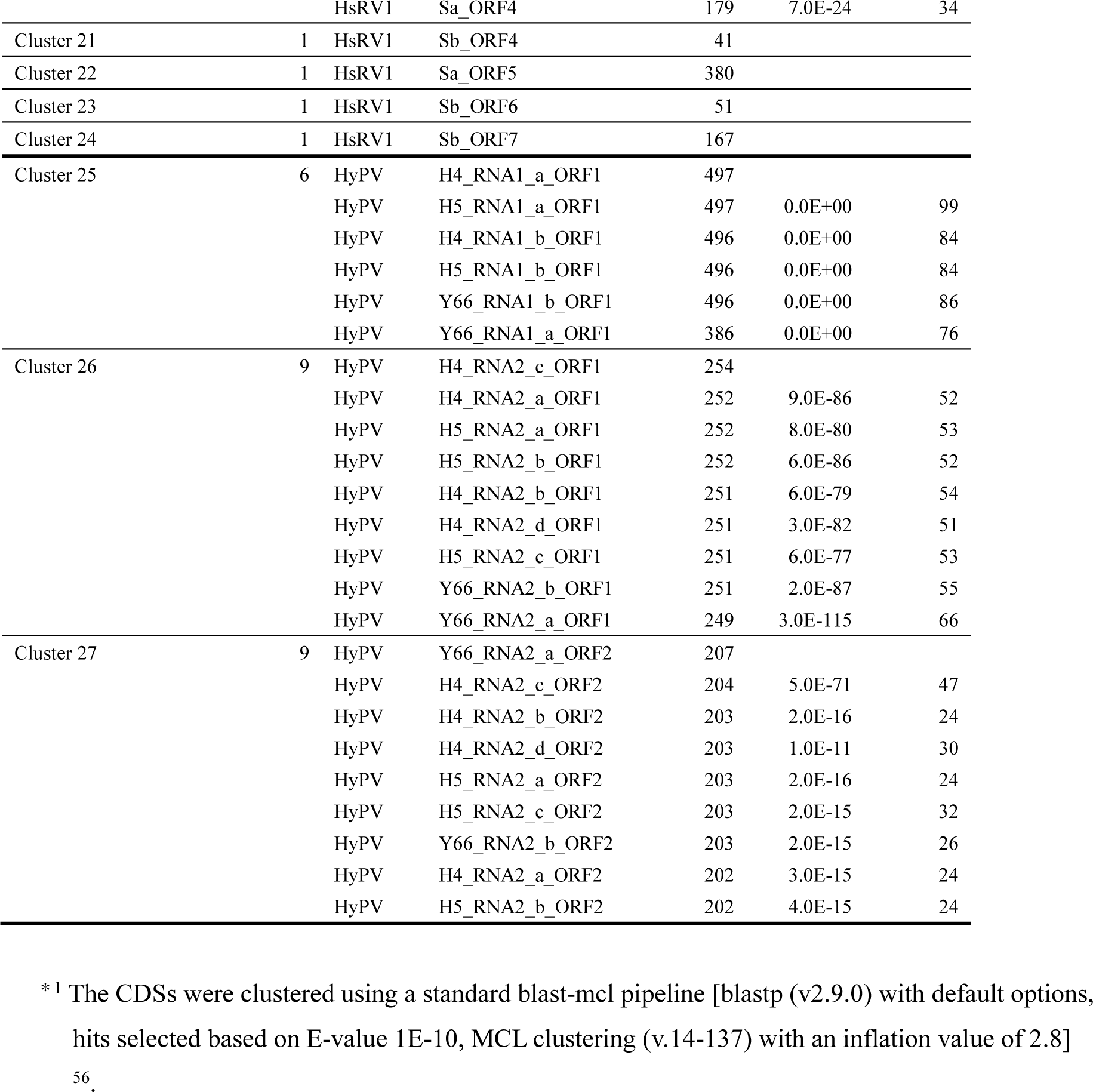
Result of CDS clustering using a standard blast-mcl pipeline.

**Table S4:**
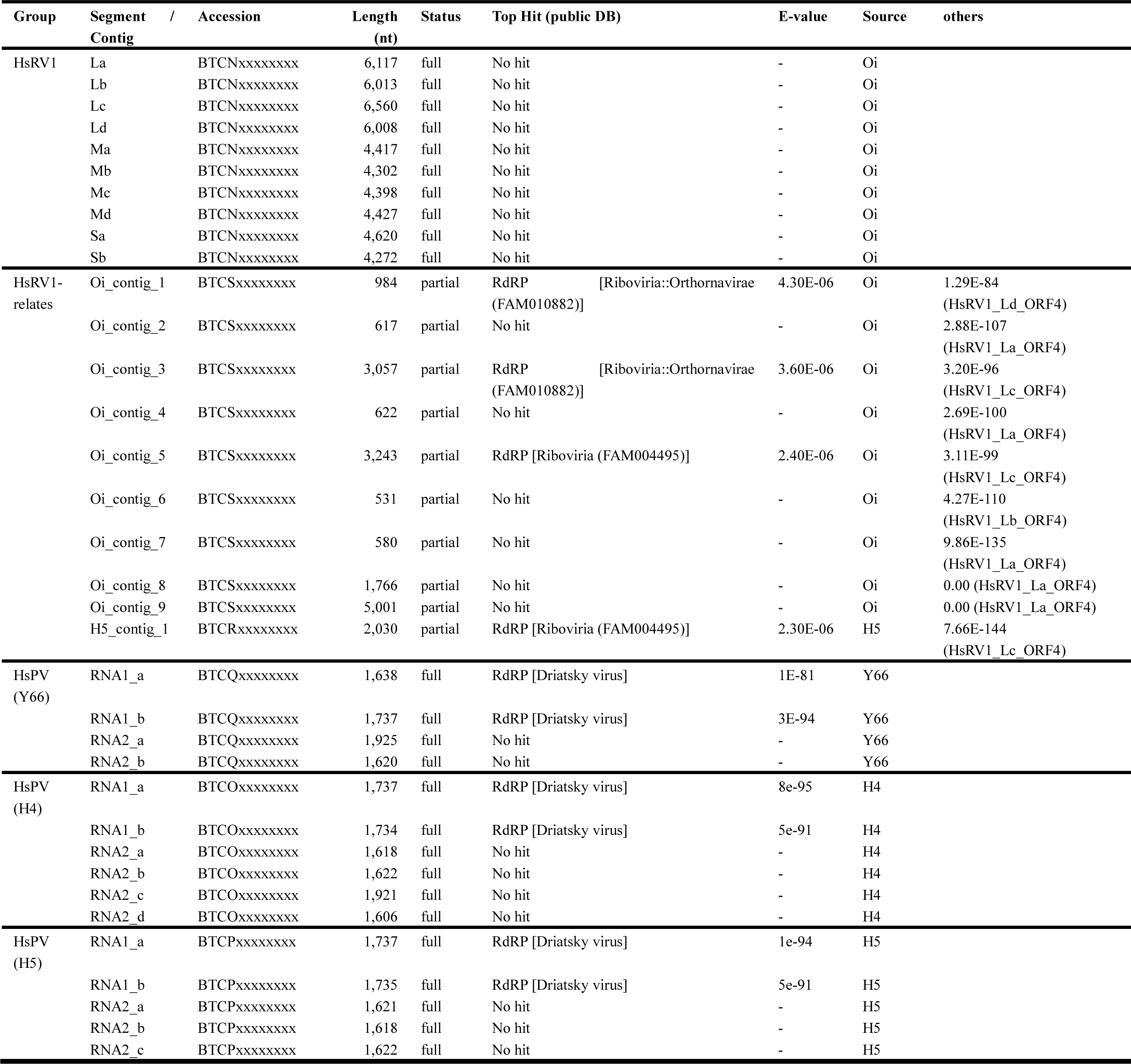
RNA virus and virus-like genomes identified in this study.

**Table S5:**
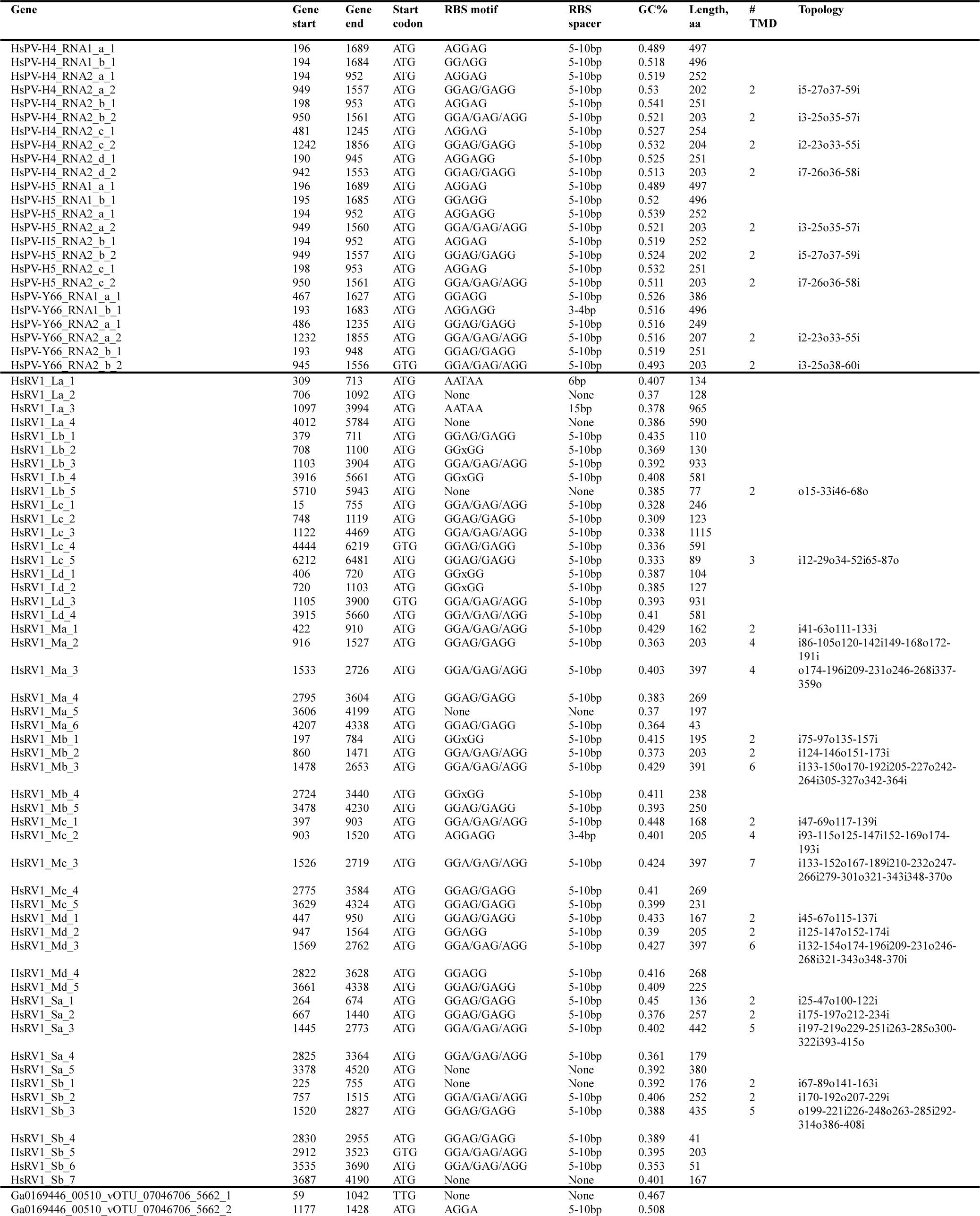
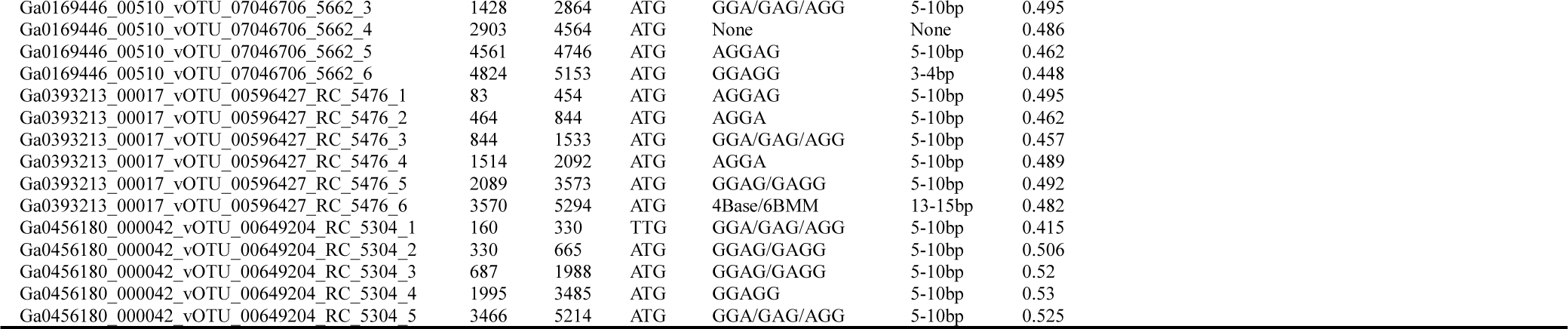
Detected RBS motif.

**Table S6:**
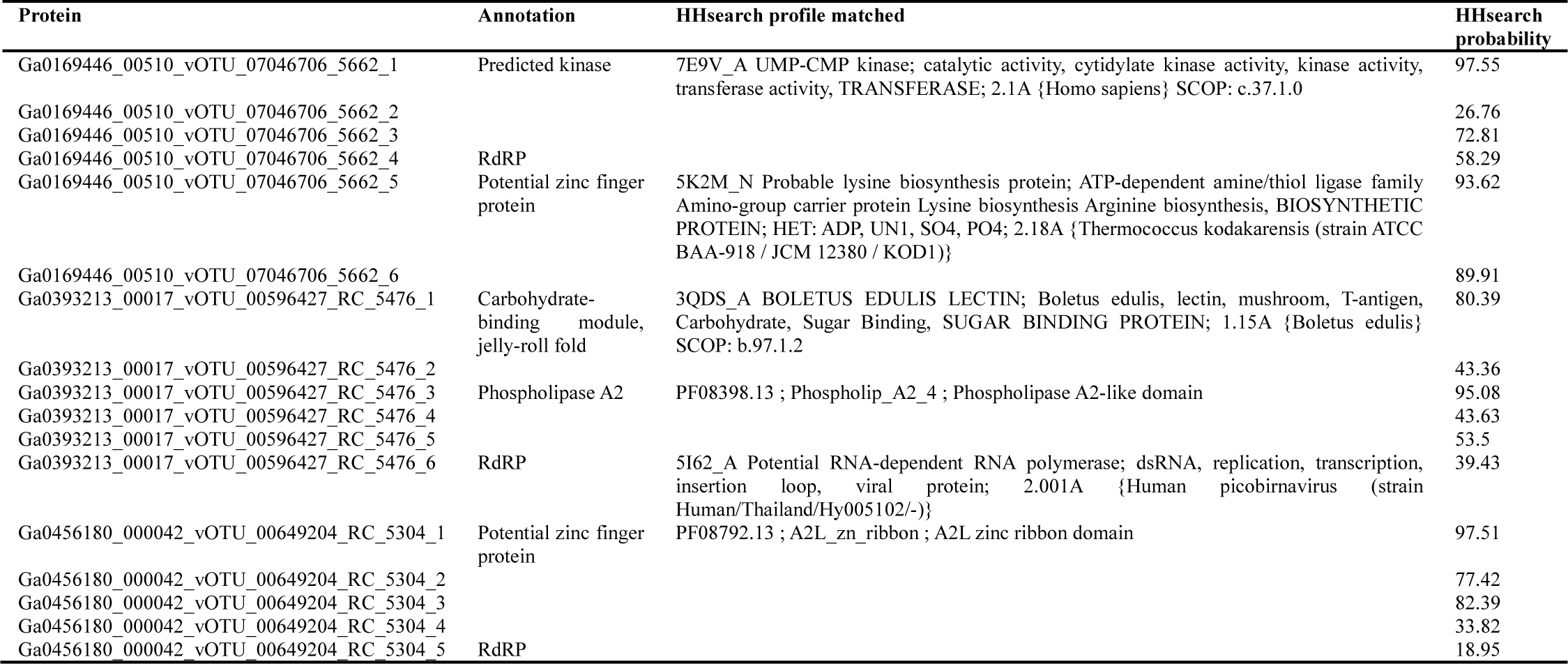
HHsearch hits for the IMG/VR virus proteins.

**Figure S1.**
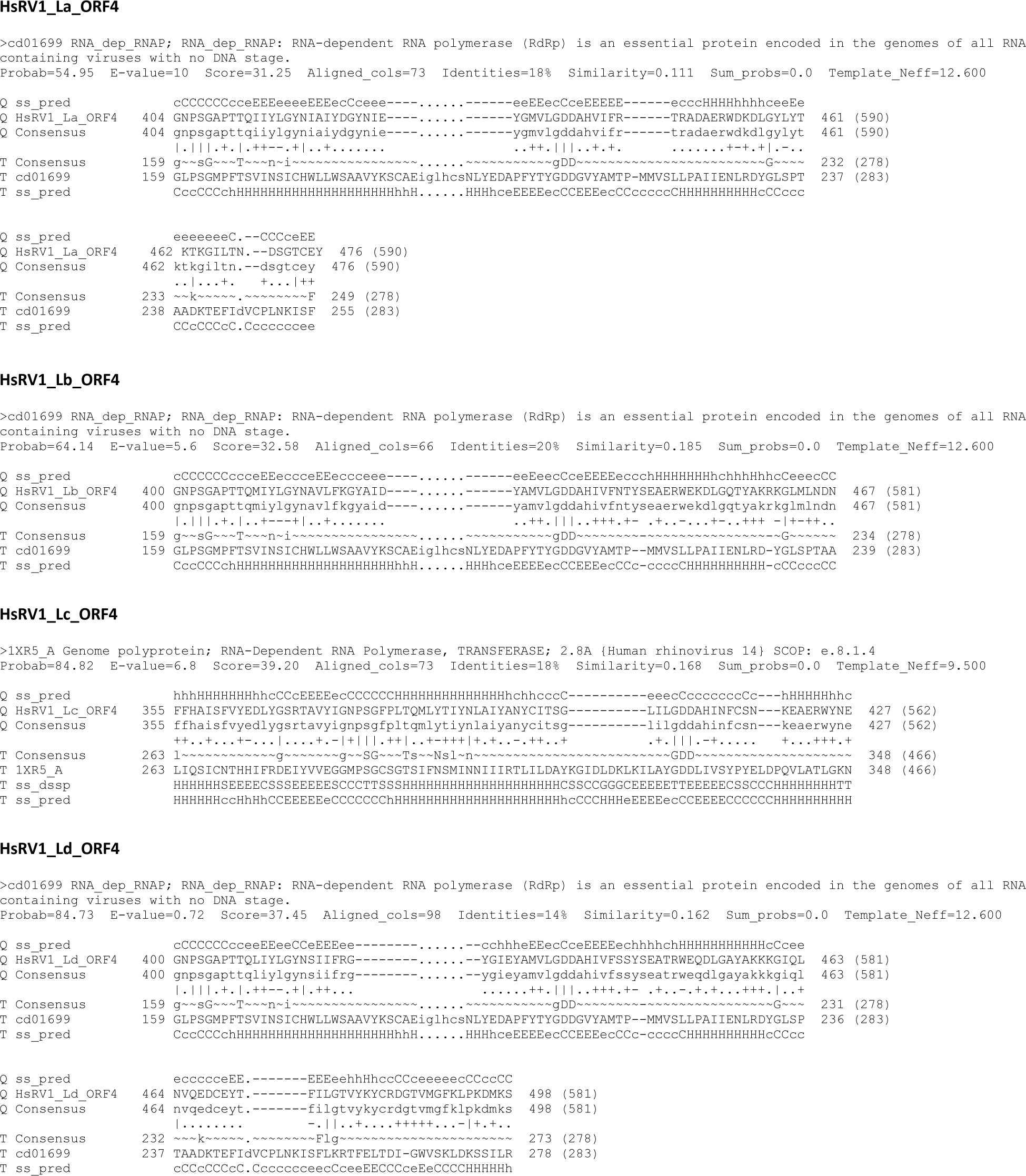
Results of the HHsearch analysis queried with the putative ORF4 protein sequences from HsRV1 virus strains. H(h), α-helix; E(e), β-strand; C(c), coil.

**Figure S2.**
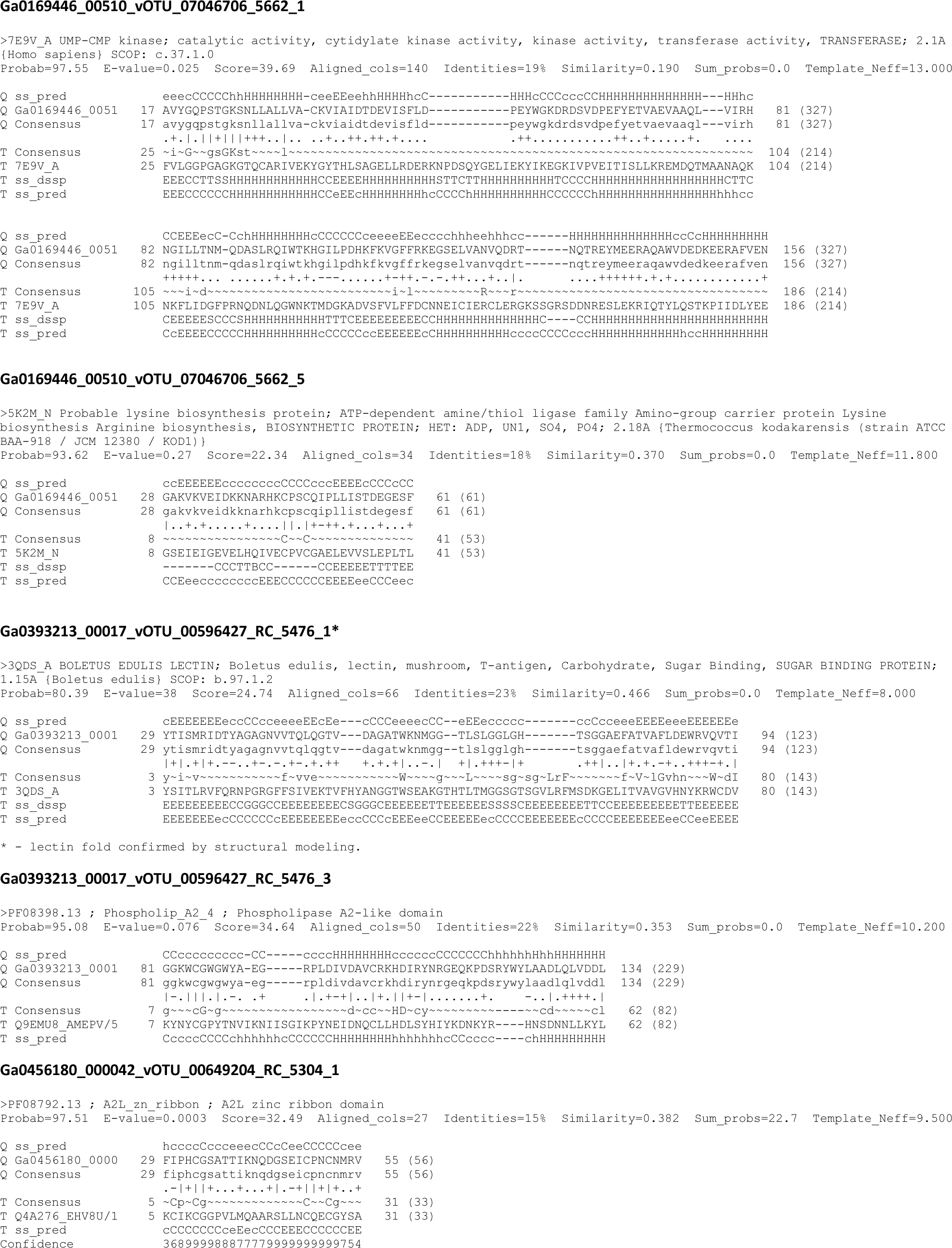
Results of the HHsearch analysis queried with the indicated protein sequences encoded by HsRV-like viruses from moderate ecosystems. H(h), α-helix; E(e), β-strand; C(c), coil.

**Fig. S3:**
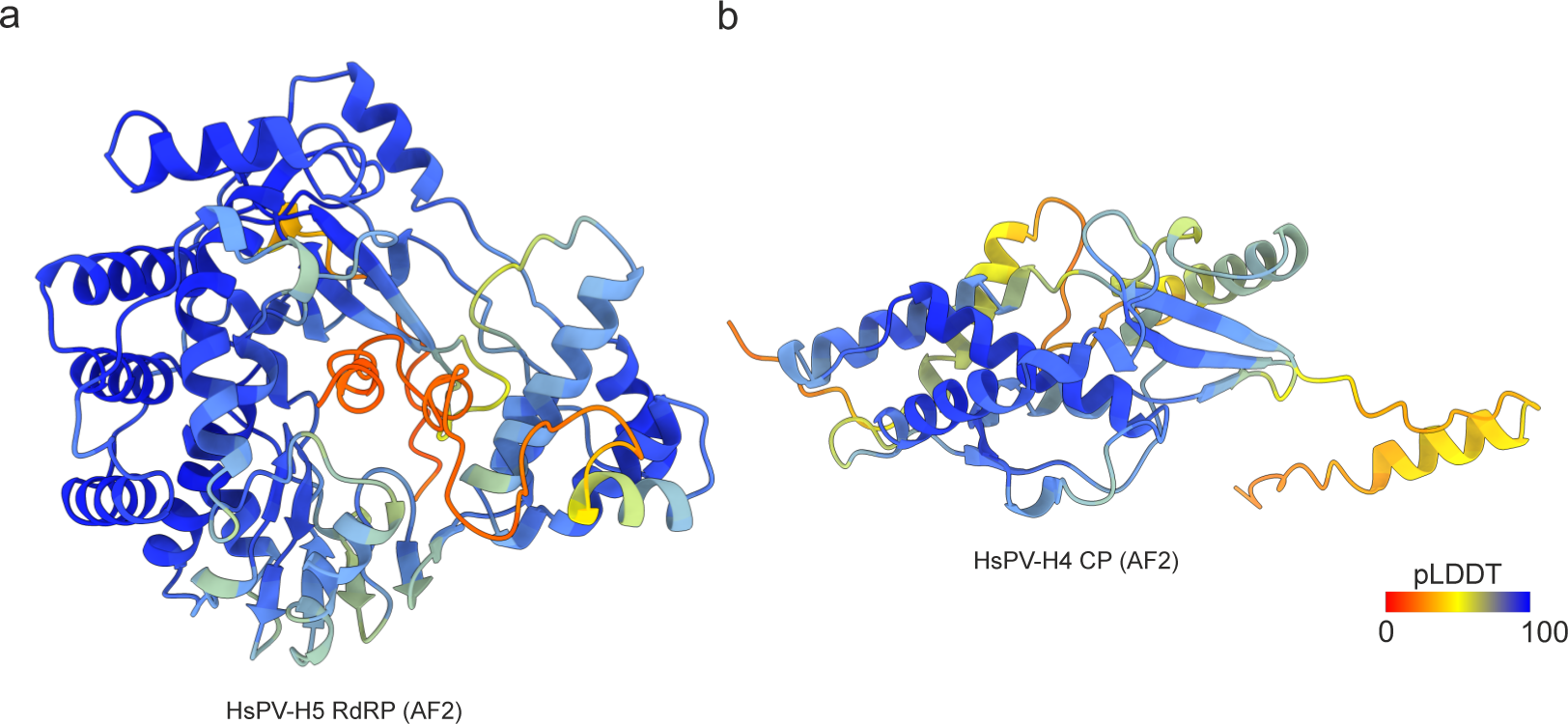
pLDDT scores of HsPV RdRP and CP. Quality assessment of the AF2 model of the HsPV **a,** RdRP and **b,** CP. The structural model is colored based on the pLDDT scores, with the color key shown at the bottom right corner.

**Fig. S4:**
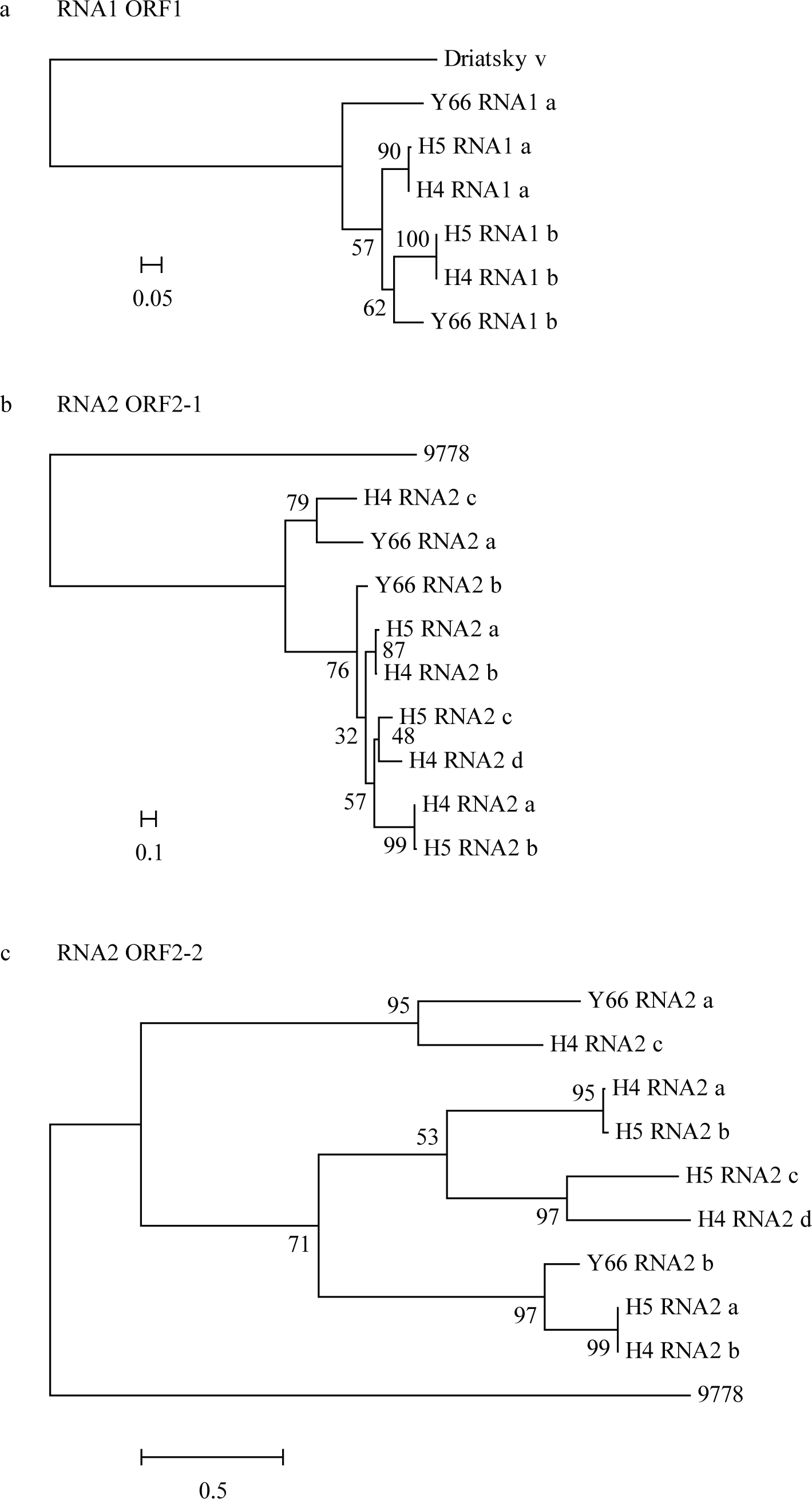
HsPV phylogeny. Maximum-likelihood trees of each ORF encoded by HsPVs and related sequences. Numbers indicate the percentage bootstrap support from 1,000 RAxML bootstrap replicates. We used RAxML with the **a,** LG+G+I+F model for ORF1, **b,** LG+G model for ORF2-1 and **c,** LG+G+I+F model for ORF2-2

