## Supplemental Information for "Distinct groups of RNA viruses associated with thermoacidophilic bacteria"

### Sampling sites

The sites H4 and H5 in Hayashida hot spring were located at a natural venting site on the slope of the valley line at the southwestern foot of a caldera lake called Onami Pond in the Kirishima Volcanic complex that is one of the most active volcanic sites in Japan. The sites T1-4 in Tearai area and Y66, Y80, and Y86 in Yunoike area were located at fumerole zone at the western foot of the same caldera pond. The site Oi was located at fumerole zone western foot of the Unsen volcano, which is a volcano that erupted in a devastating eruption that lasted from November 1990 to February 1995 [1]. The site Ob was in Obama hot spring area, which is located ~10 km apart coastal area from the site Oi (Fig. S4).

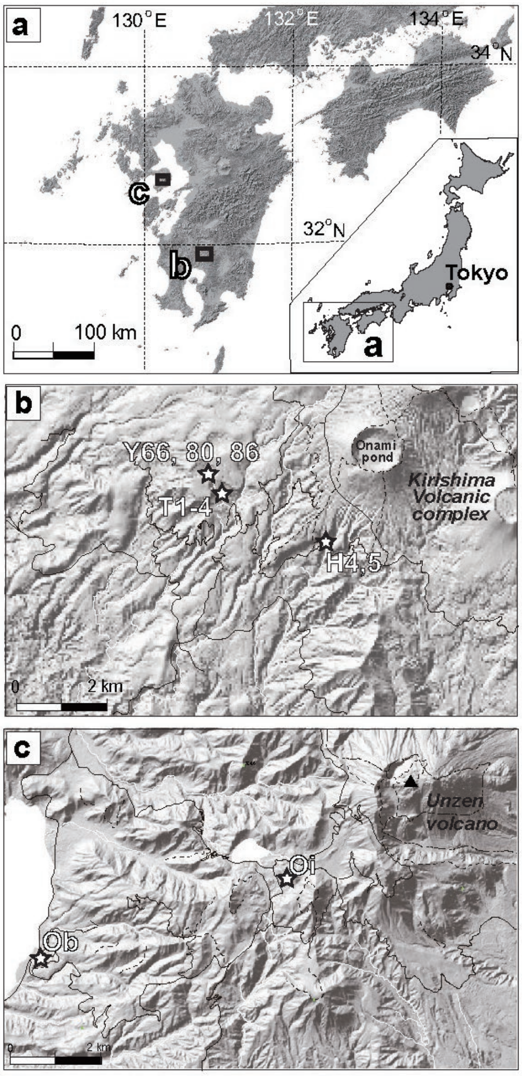

**Figure S4. Locations of the sampling points. a,** Samples were collected from Kyushu, southwestern Japan, where an area with many active volcanoes. **b,** Three sites (H: Hayahida hot spring, T; Tearai area, and Y; Yunoike area), were located on the west side of the Kirishima Volcanic complex. **c,** The sites Oi was located at a fumerole zone western foot of the Unsen volcano. The site Ob differs from the other sites in that it was located along the coast, relatively far from an active volcano. The maps were based on the digital elevation topographic map published by Geospatial Information Authority of Japan（http://maps.gsi.go.jp/）.

### Chemistry of hot spring water

The chemical composition of hot spring water was measured. The major cation ($Ca^{2+}$, $Mg^{2+}$, $Na^{+}$, $K^{+}$) and anion ($Cl^{-}$, $SO_4^{2-}$) were analyzed using ion-chromatography (Gulliver; Jasco International Co., Ltd., Tokyo, Japan) after 21 times dilution of the samples with distilled water. $SiO_2$ was measured by the molybdenum yellow method with a spectrophotometer. δD and $\delta^{18}O$ were measured from filtered (0.2-μm-pore-size) and non-diluted water samples by using a liquid water isotope analyzer (Los Gatos Research, Inc.). Measurement errors are within 5% for $Ca^{2+}$, $Mg^{2+}$, $SO_4^{2-}$, and $SiO_2$, 10% for the other ions, ±1.1‰ for δD, and±0.1‰ for $\delta^{18}O$. Table S6 shows the measurement results of the water samples.

Table S6. Chemical and isotopic composition of the fluid samples at the 11 samples.

| Code | $Ca^{2+}$ (mg/L) | $Mg^{2+}$ (mg/L) | $Na^{+}$ (mg/L) | $K^{+}$ (mg/L) | $Cl^{-}$ (mg/L) | $SO_4^{2-}$ (mg/L) | SiO2 (mg/L) | δDH2O (‰ SMOW) | δ18OH2O (‰ SMOW) |
|---|---|---|---|---|---|---|---|---|---|
| H4 | 181 | 7 | 77 | 112 | 15 | 33 | 142 | -49.2 | -7.1 |
| H5 | 195 | 7 | 80 | 108 | 15 | 33 | 141 | -48.1 | -6.9 |
| T1 | 238 | 8 | 995 | 106 | 419 | 111 | 150 | -35.2 | -3.3 |
| T2 | 401 | 16 | 81 | 132 | 10 | 602 | 271 | -25.1 | -1.5 |
| T3 | 265 | 10 | 77 | 140 | 10 | 170 | 188 | -31.0 | -3.0 |
| T4 | 223 | 10 | 62 | 128 | 10 | 304 | 183 | -33.5 | -0.7 |

| | | | | | | | | | |
|---|---|---|---|---|---|---|---|---|---|
| Y66 | 195 | 8 | 53 | 117 | 10 | 31 | 92 | -40.6 | -4.7 |
| Y80 | 202 | 8 | 53 | 118 | 10 | 45 | 98 | -18.8 | 3.8 |
| Y86 | 185 | 8 | 49 | 97 | 30 | 114 | 187 | -41.9 | -4.3 |
| Oi | 26 | 4 | 22 | 9 | 1 | 431 | 119 | -30.6 | -4.4 |
| Ob | 113 | 139 | 2739 | 310 | 1671 | 131 | 133 | -31.8 | -3.8 |

**Contig sequence**

>SRR7239362_trimmed_contig_9778 Average coverage: 49.10

GTTCTGTCTAAACACTAGCCTTCCACTTAGTGGTGATAGCTTAATTAGTTAG
GTAAACTAAAGCTGACCGTCCTGTCTCTGGGCCCGCTGGTTGAGGTAACACC
CGTATTTAGGACAACGTATGTCTGGCCTTCGGGCAAGTCGGCATTCCGCCGG
CGATTTCTCGAATCATGGACGAATTCGAGTCATTCCGTCAGTCCCGTCCTTA
TAGGCAGCATAGTGCGCCTGAGGTCTCAGTACGTAAACTGTTTACTTTGGAG
TCTTTATGATTAAACATAATACTCAAGCTGTCAGTGATGATAGCATTGAACC
TGTTGCCCGTCAGTACGGTCTTCAACTGATGGCACAGGGGGTGGCAACACT
CTCCCAATGTGACGTCTTCGTTCGCCCCATCCCTGGTATCCAATGTGTTGTTG
GTGCTTGGGCTACTCTTACGAAGTTCAAACGAGCTGCTCAAGTTGTTTCCGA
GGAACGCCTCAAAGAACACTTGAACATCGTCACTGCTTTACGCGTCCTTCAA
GTTCGTGGTGAGATTGATGACCCGCGTATGGATGTTCGTCGCTGTGTCTATC
CCTCGATCCTCCGCCCGGTGTTCCGTGCGATTGGTGATGTGATGGATGAGAG
CGTGAGCCTCGATCTTCGCGTTAATCTTTCCGATGAGCTTGTGGAAGTCATT
AAAGGGTATGACTTCGCAAACTGGCGTGACGACAACCAAATGATTCAGATC
GCTTTAATGCAATCTGGTATCTCCTGCGCTACCTGCCTACCGCCTGAGGTCG
ACGGGAATCGTCAGGTGTTGACCATGATCGTTAAGGAGGTCGAGGGTCAGG
CCCGAGCCGAGGGTCAGGTCGTCGGTTTTGATCGTTCAGCGAATCCTGGTGA
GGTTCTGGTCGGAGCCATTTTGGGCCACCGTTTAGAAGATCCTTCCATCTTC
GGTGCTCCGCGTGTCTCATACCAAAACGTCGGGTACTTCGAAGACCAGTTCT
ATAACTTGGTTAACGCTGAAGTTCTCAACGCCCGTTCTAAGTAGTGATGAAA
GAGAGGTCCAGCGATGAATCTATCCTTATTCTTTGTAGCATTTGTAATCTTCT
GGTTACTATCTGTGGTCATATGCTTCATGGTAATTGGGTTCGTTAATGGCCT
CTCGTTCGGCTTCGCTTTCCGTTACCTGACCACAGGTGATGGCGCGTTGCTC

GTCACCCTTACAGCCAGTGTATGTGAGATATTCTACTTACTATTCTGGCTTC
ACTGCGTGCTAGATCAGCGTTTGGCAGTCGAGGATGTTTCTCGTCAAGTCGA
GAAGATACGCACGAGTATCAACAAGATTAGTCGAGACGATTTGTCAAAGCT
CAATGCCGTTGTGGATGGCGTTAATTCGCTTAATACTGAGGTCCTTCGTCTG
GAGAAGGGGCTTGATGAAGCCTGTGATGTTATGGAAAGGAACATCCCTAGG
TTAAGTCAGTCTGCTGACGAGATCGTTCGGTTCCACAGTGCATTGGACTCCC
ATGGTGAGGAGTTGAGAGAGATGTTCGTGAATACACTCGCATCTCGATATC
ACGAGGTCCTGACGGAAACCTTCAGTGAGACCAAACTAGCACAGATACGTG
GGTTTGTATCAAGGCAGTCGTTTAAGATTCTCACGTTGCATTTGCCATTCTGT
GAGTATTTCGACTTGTTCACTCCCGAGCAGAAGAAGTTGTACTTCGACTGTA
TGCGTAGCGGCGATTACAGACGCCTGTCGAAGAGCCTGACGAAGGAACAGA
GAGACGCTGTAAGTAAGGGTTAGGTCTCCCTTGCGGAGGGGGGG

**Metagenomic sequencing and data processing**

Five metagenomic libraries were constructed and analyzed as described before [2] (Table S7). In brief, DNA was extracted from cells collected on a portion of the 0.2-μm-pore-size filters corresponding to approximately 0.25–2.5 L of H4, H5 and Oi samples, that potential complete RNA virus genomes were identified, using DNeasy PowerSoil Kit (QIAGEN). Covaris M220 (Woburn, MA, USA) was used for physical DNA fragmentation using the conditions described below to obtain a peak fragment size: 400 bp; Peak Power: 75.0, Duty Factor: 15.0, Cycles/Burst: 200, and Time: 60 s. Then, shotgun metagenomic libraries were constructed using KAPA Hyper Prep Kit. The metagenomic sequence libraries were analyzed using the Illumina MiSeq platform with a 2×300-bp read length.

Sequence reads were quality filtered using Trimmomatic v0.35 with the option "LEADING:20 TRAILING:20 MINLEN:60". The quality-controlled reads were then assembled in a sample-by-sample manner using MEGAHIT v1.1.4 with the default setting. Only the long contigs (≥1kb) were retained for further analyses. CRISPR regions were identified using MinCED v0.4.2 with the default setting. The identified CRISPR spacers (n=919) were assembled as a database for the identification of potential virus-host interactions. The 25 virus genome segments were used as a query for sequence similarity search against the CRISPR spacer database. The similarity search was performed using BLASTn with the options "-word_size 7 -evalue 1e-3 -dbsize 100000000" and no sequence met the threshold.

Table S7. Hot spring metagenomes and CRISPRs

| sample | read bp | assembly size (≥1kb) | number of contigs | CRISPR region | CRISPR spacer |
|---|---|---|---|---|---|
| H4 | 631873847 | 17009520 | 8982 | 19 | 66 |
| H5 | 555047200 | 4338329 | 1735 | 15 | 100 |
| Oi | 694063525 | 16906462 | 11827 | 62 | 207 |
| Y66 | 622940918 | 18786560 | 11513 | 56 | 347 |
